# Representation, Alignment, and Generation: A Comprehensive Survey of Foundation Models for Non-Invasive Brain Decoding

**DOI:** 10.64898/2025.11.30.691403

**Authors:** Yifan Wang, Shaonan Wang, Wenhao Cai, George Ford, Yang Cui, Yunhao Zhang, Changde Du, Cunhang Fan, Dongyang Li, Hongpeng Zhou, Hongyu Zhang, Jixing Li, Quanying Liu, Wei Huang, Yizhuo Lu, Zijiao Chen, Jingyuan Sun

## Abstract

The ability to decode human thoughts, intentions, and perceptions directly from non-invasive brain recordings holds transformative potential for healthcare, communication, and human-computer interaction. However, translating the safety and scalability of methods like Functional Magnetic Resonance Imaging (fMRI), Electroencephalography (EEG), and Magnetoencephalography (MEG) into real-world utility has traditionally been hindered by low signal-to-noise ratios, limited spatial-temporal resolution, and the difficulty to collect large-scale high-quality data from an individual user. Recently, the emergence of Foundation Models (FMs)—large-scale, pre-trained architectures—has expanded the feasible operating region of non-invasive brain decoding under controlled protocols, primarily through representation learning, neuro-semantic alignment, and strong generative priors. However, evidence for robust cross-subject and real-world deployment remains uneven, as many results are still demonstrated on limited cohorts or highly controlled settings. This survey provides a comprehensive overview of how FMs are redefining the boundaries of non-invasive brain decoding. We propose a unified methodological framework that synthesizes recent advancements into a coherent process: extracting robust, transferable representations from noisy neural signals; aligning these signals with the rich semantic spaces of pre-trained vision and language models; and leveraging powerful conditional generative priors to reconstruct high-fidelity outputs. We systematically review state-of-the-art applications across three key domains: visual reconstruction, language and speech decoding, and auditory processing. Furthermore, we critically examine the persisting challenges of computational efficiency, cross-subject generalization, and privacy governance. By mapping the current landscape and identifying key gaps, this work outlines a strategic research agenda aimed at transitioning FM-driven neurotechnology from laboratory proofs-of-concept to reliable, real-world applications.

## 1 Introduction

The enduring quest to understand the human brain and translate its intricate activity into meaningful outputs represents a central challenge in neuroscience and engineering (da Silva 2013). Decoding brain signals—interpreting neural activity to infer cognitive states, intentions, or perceptual experiences—has garnered significant interest due to its profound potential. Laboratory research has yielded remarkable progress, demonstrating the feasibility of interpreting neural signals to control prosthetic limbs, enable communication for individuals with severe paralysis, and even reconstruct perceived visual stimuli from brain activity patterns (Naselaris et al. 2009; Kay et al. 2008; Willett et al. 2021b). These achievements underscore the transformative possibilities, spanning healthcare innovations for neurological disorders, novel communication channels, and enhanced human-computer interaction paradigms (Pais-Vieira et al. 2013).

To bring these technologies from the laboratory to the general public, researchers have increasingly focused on non-invasive brain decoding techniques, such as Electroencephalography (EEG), Magnetoencephalography (MEG), Functional Magnetic Resonance Imaging (fMRI), and Functional Near-Infrared Spectroscopy (fNIRS) (da Silva 2013; Buzśaki et al. 2012; Logothetis 2008; Ferrari and Quaresima 2012). Unlike invasive methods, these techniques acquire neural signals from outside the body without surgical intervention, thereby avoiding associated health risks (Rossi et al. 2021). Their safety and scalability make them a promising foundation for widespread use. However, non-invasive methods face significant inherent challenges: they typically suffer from lower signal-to-noise ratios (SNR) and poorer spatial or temporal resolution compared to invasive implants. Furthermore, high-quality physiological data is relatively scarce, and data annotation is costly. These factors have significantly hindered the development of robust, generalizable decoding systems (Abiri et al. 2019; Rubin et al. 2017).

Historically, brain decoding largely relied on linear models or task-specific deep networks. While effective in controlled paradigms, the performance of these models often collapses when applied to naturalistic environments or when generalizing across different subjects. In recent years, however, a paradigm shift in artificial intelligence (AI)—driven by the advent of Foundation Models (FMs)—has presented new opportunities to overcome these limitations (Radford et al. 2021; Gong et al. 2024). FMs are large-scale machine learning models pre-trained on vast and diverse datasets, often using self-supervised learning, which allows them to adapt to a wide range of downstream tasks (Radford et al. 2021; Devlin et al. 2019; Gong et al. 2024). Architectures such as the Transformer (Devlin et al. 2019) have enabled models like GPT (Radford et al. 2019) in language and CLIP (Radford et al. 2021) in vision to capture intricate semantic patterns. By leveraging these capabilities for representation learning, feature extraction, and cross-modal alignment, FMs have significantly improved the fidelity and semantic accuracy of reconstructions from non-invasive brain recordings, opening unprecedented avenues for neurotechnology (Gong et al. 2024).

Although reviews on brain decoding are extensive, existing literature exhibits key shortcomings that this survey aims to address. Previous surveys have often been limited in scope or outdated relative to the rapid pace of AI development. For instance, reviews on EEG applications (Värbu et al. 2022) or encoding/decoding (Xu et al. 2021; Abiri et al. 2019) often focus solely on specific modalities or emphasize algorithmic lists without unifying visual, linguistic, and multimodal fusion. Similarly, surveys on natural image reconstruction (Rakhimberdina et al. 2021) and fMRI decoding (Du et al. 2022) highlight limitations like data scarcity and hemodynamic delays but do not fully explore how modern FMs resolve these issues through ”neuro-semantic alignment” and conditional generation. A unified framework that connects the three major task families—visual (image/video), language/speech, and cross-modal fusion—remains lacking.

This paper addresses this gap by proposing a comprehensive methodological framework: using transferable representations as a foundation to align brain signals with the semantic or acoustic latent space of large models. We review how ”weak, sparse, and delayed” neural measurements can be transformed into readable images, text, and speech by performing conditional generation or retrieval within this aligned space. From this perspective, we systematically review advancements in image and video reconstruction, language and speech decoding, and auditory attention. We map datasets and evaluation metrics to this framework, emphasizing practical challenges such as cross-subject generalization, inference efficiency, and privacy governance.

The remainder of this survey is organized as follows. Section 2 establishes the theoretical foundations that motivate aligning computational models with biological neural processing, including neural coding, representational geometry, and related perspectives. Section 3 surveys the non-invasive decoding landscape by introducing major modalities and their constraints, and further summarizes commonly used datasets and benchmarks (Figure 1; Tables 1 and 2). Section 4 reviews core concepts and representative architectures of foundation models relevant to brain decoding (Table 3). Section 5 presents foundation-model–based decoding advances across vision, language/speech, and auditory attention/audio, guided by an overview timeline (Figure 2) and task-specific schematics (Figures 3 and 4), with datasets and reported evaluation metrics consolidated in Tables 4, 5 and a dedicated discussion on metric interpretation and compatibility. Section 6 synthesizes cross-domain trends and recurring methodological patterns. Section 7 discusses persisting challenges and future horizons, including validation regimes and reproducibility considerations, applicability boundaries contrasting FM-based and classical pipelines (Table 6), and a prioritized research roadmap (Table 7). Finally, Section 8 concludes with key takeaways and requirements for translating FM-driven non-invasive brain decoding from controlled demonstrations to reliable real-world applications.

**Fig. 1.**
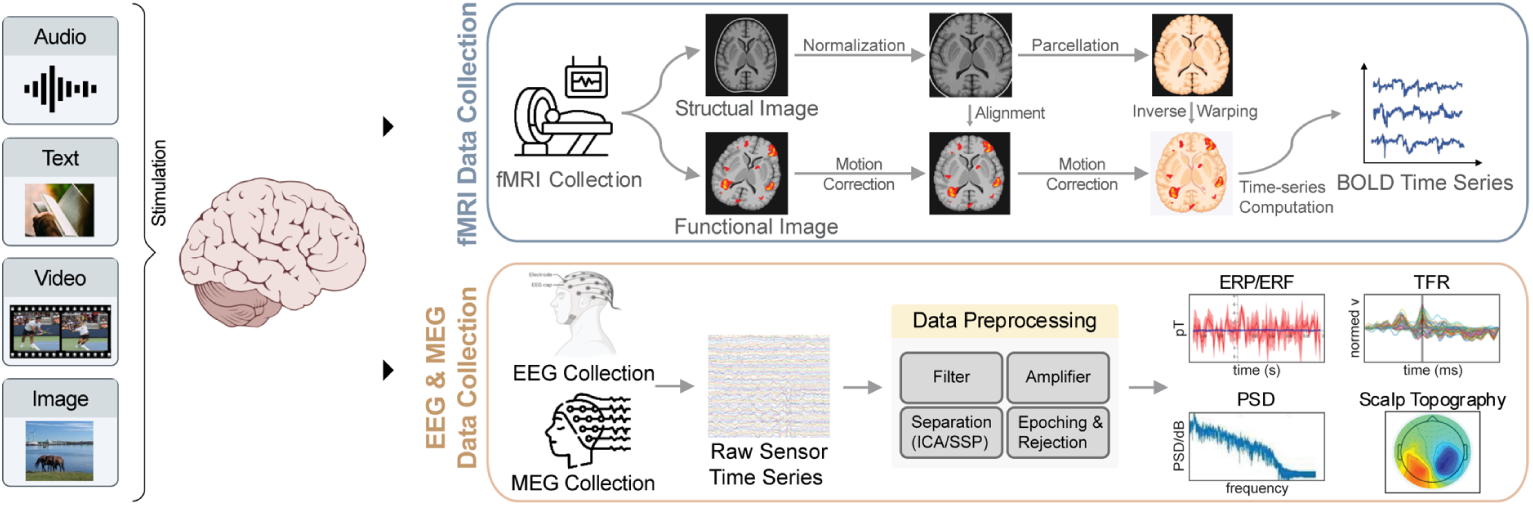
Non-invasive data collection and preprocessing across fMRI, EEG and MEG. The diagram highlights typical stimuli, acquisition hardware, core preprocessing steps (e.g., normalization/alignment/motion correction for fMRI; filtering/ICA/epoching for EEG/MEG), and the resulting analysis-ready outputs (BOLD time series, ERPs/TFR/PSD/topographies).

**Fig. 2.**
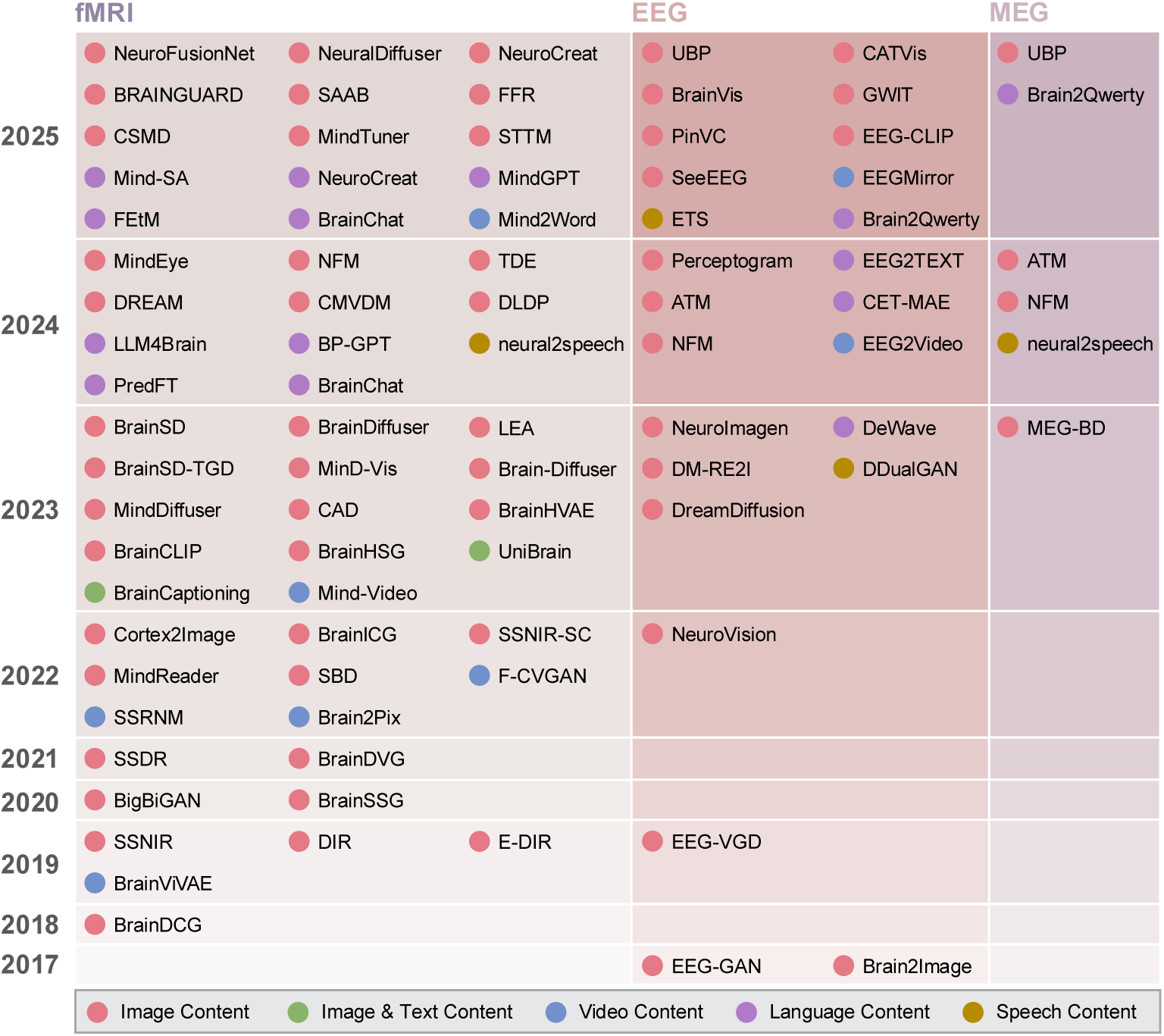
Chronological map of representative non-invasive brain-decoding works (2017–2025) across modalities (fMRI/EEG/MEG). Colors denote output content types (image, video, language, speech). This overview situates subsequent method sections and helps cross-reference modality-specific trends.

**Fig. 3.**
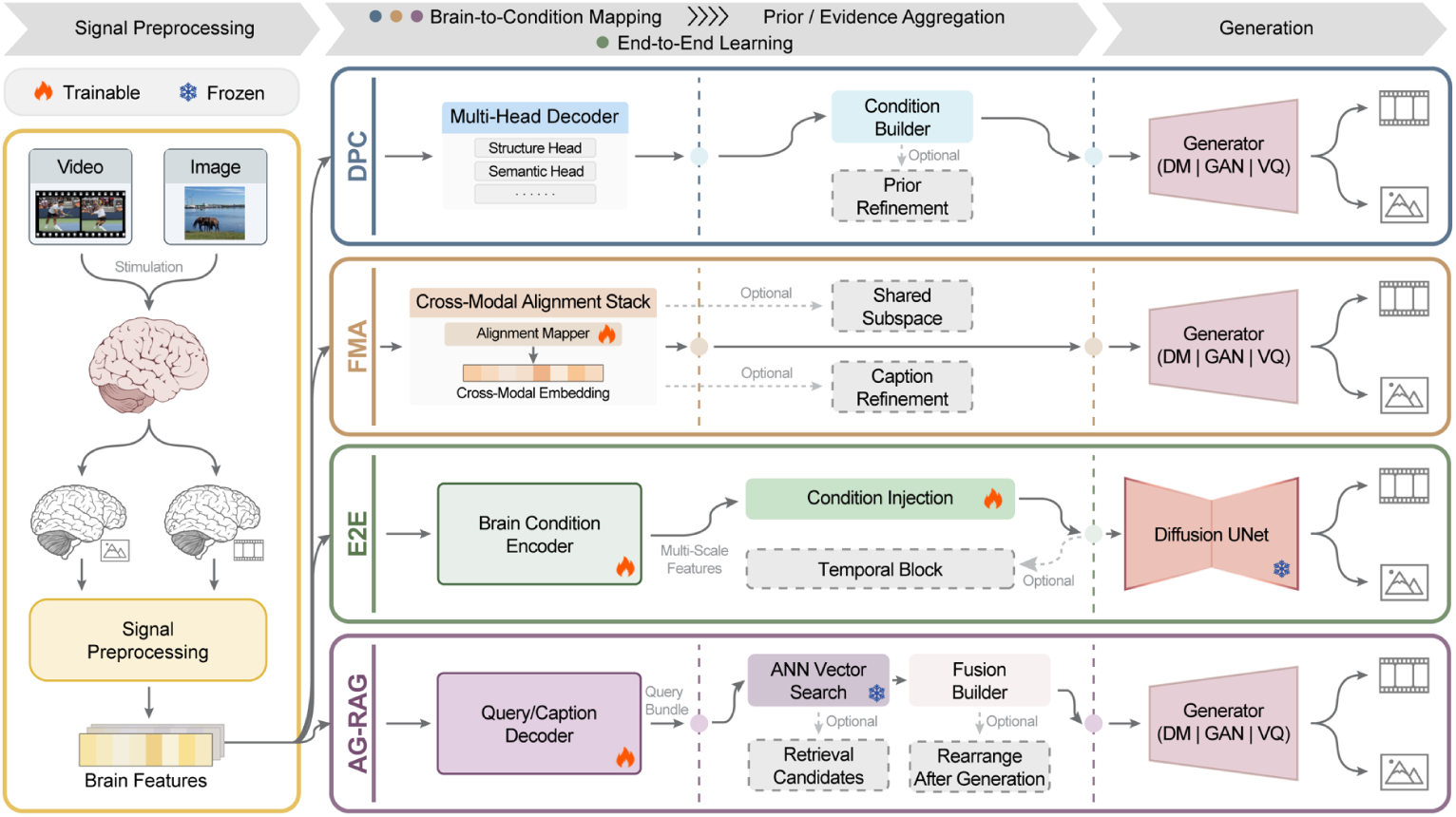
Decoding Visual Experiences Flowchart. This flowchart shows the different types of methods for images and video decoding tasks. (1)DPC (Decoded-Prior Conditioning): Multi-head decoders predict structural (VAE latent, depth/edges) and semantic (CLIP/caption) priors, fused and fed to a generator (DM/GAN/VQ); dashed boxes denote prior refinement/re-ranking options. (2) FMA (Foundation Model Adaptation (Alignment)): Brain features are aligned to multimodal embeddings (CLIP/text), optionally with shared-subspace learning and caption refinement, then passed to a generator (DM/GAN/VQ) and diffusion. (3) E2E (End-to-End Decoding): A learned brain condition encoder injects multi-scale features into a largely frozen diffusion UNet; an optional temporal block supports video. (4)AG-RAG (Augmented Generation via Retrieval-Augmented Generation): A query/caption decoder drives ANN search over a multimodal index; retrieved candidates supply latent init, geometric controls, and prompts to the generator (DM/GAN/VQ/Retrieve-only), with optional post-generation re-ranking.

**Fig. 4.**
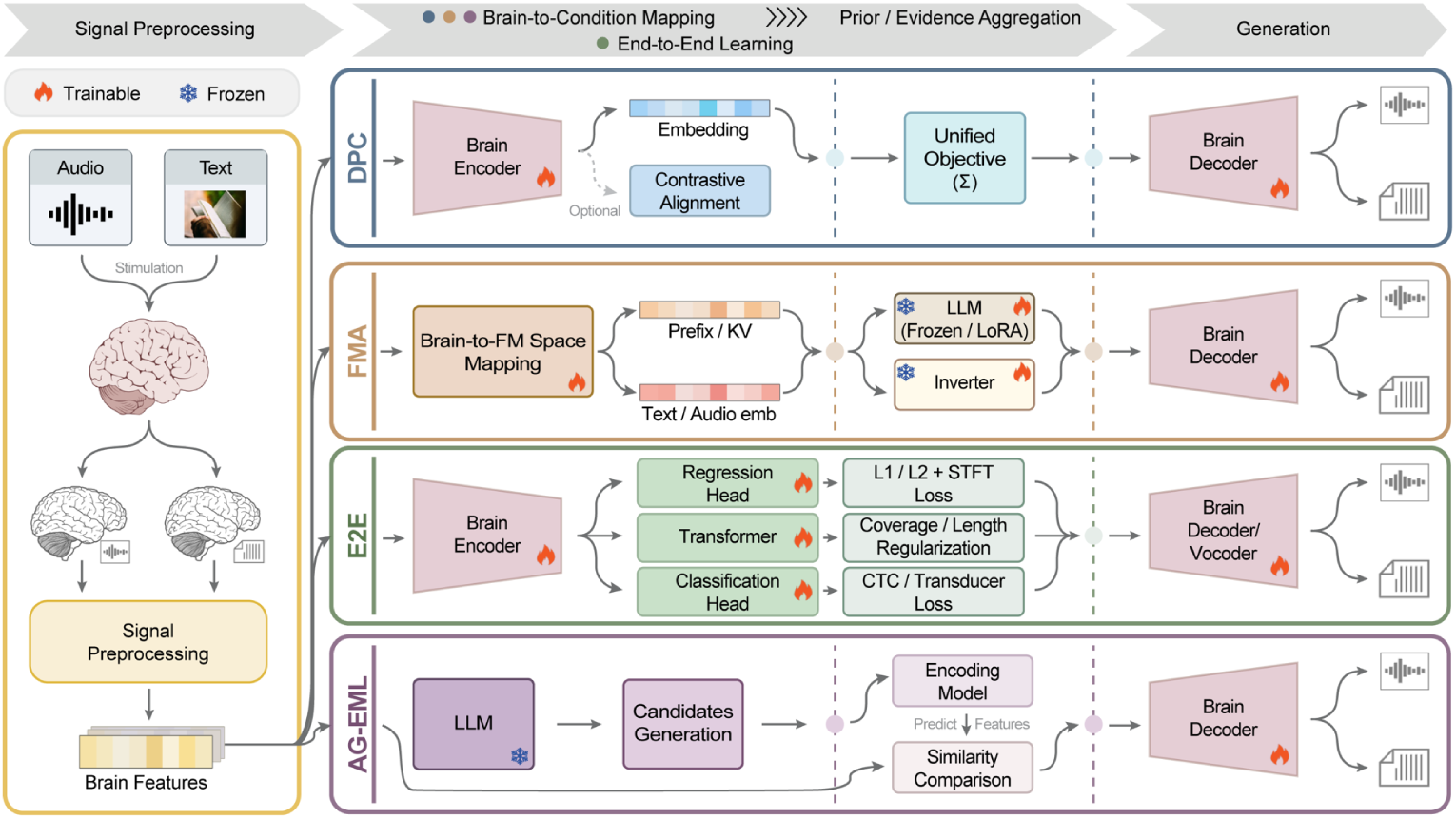
Decoding Language and Speech Flowchart. This flowchart shows the different types of methods for language and speech decoding tasks. (1) DPC (Decoded-Prior Conditioning): Preprocessed brain signals are encoded by a Brain Encoder and decoded by a Unified Decoder (text/speech) under a Regularized Multitask Objective (CE, spectral losses, alignment, regularization); the dashed Contrastive Alignment denotes an optional brain–semantic alignment. (2) FMA (Foundation Model Adaptation): A Mapping (Brain→FM Space) projects brain features into the foundation-model space, followed by two routes: (i) injecting context vectors/keys/pre-fixes to a frozen or lightly fine-tuned LLM; (ii) regressing to text/audio embeddings and using an Inverter to recover readable text or synthesizable speech. (3) E2E (End-to-End Decoding): After a shared Brain Encoder, three branches are used: Regression Head to acoustic features with L1/L2 + STFT and a Vocoder; Transformer seq2seq with coverage/length regularization; and Classification Head with CTC/Transducer loss for direct character/phoneme/word decoding. (4) AG-EML (Augmented Generation via Encoding-Model-in-the-Loop): A separately trained encoding model scores LLM-generated candidates against measured brain activity (NeuroScore), enabling reranking to select the final output.

**Table 1.** Comparison of Non-Invasive BCI Modalities.

| Modality | Principle | Spatial Resolution | Temporal Resolution | Invasiveness | Portability | Cost | Key Strengths | Key Weaknesses |
| --- | --- | --- | --- | --- | --- | --- | --- | --- |
| EEG | Electrical potentials from neural currents | Low (cm) | High (ms) | Non-invasive | High | Low | High temporal resolution, portable, low cost | Low spatial resolution, susceptible to artifacts/noise |
| MEG | Magnetic fields from neural currents | Moderate (mm-cm) | High (ms) | Non-invasive | Low (requires shielding) | Very High | High temporal resolution, better spatial resolution than EEG | Expensive, requires shielded room, sensitive to movement |
| fMRI | Blood-Oxygen-Level-Dependent (BOLD) signal | High (mm) | Low (seconds) | Non-invasive | Very Low (immobile) | Very High | High spatial resolution | Low temporal resolution, indirect measure, expensive, immobile, noisy |
| fNIRS | Light absorption by blood hemoglobin | Moderate (cm) | Moderate (seconds) | Non-invasive | Moderate-High | Moderate | Portable, balanced trade-off, movement tolerant | Lower spatial resolution than fMRI, limited depth penetration |

**Table 2.**
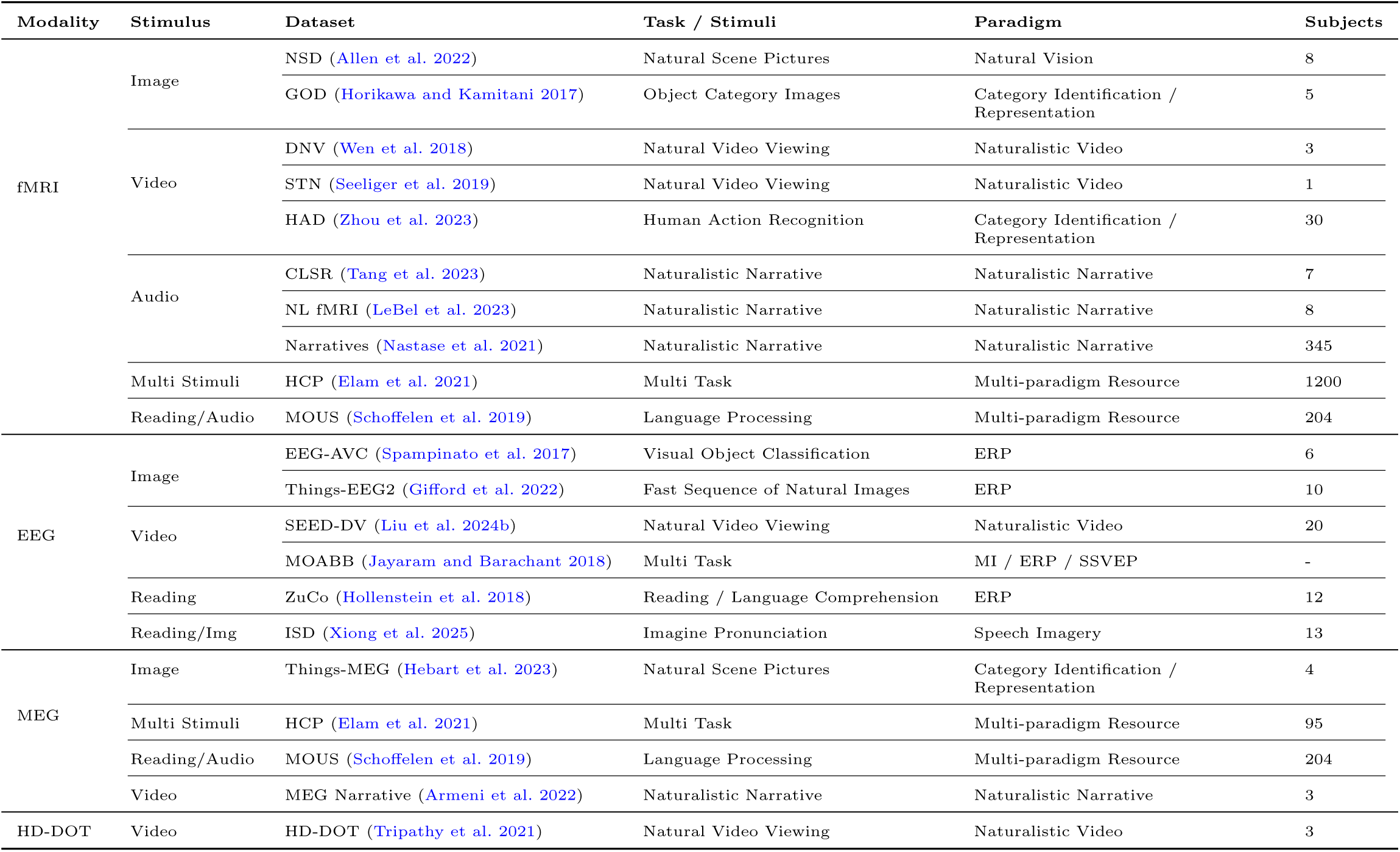
Summary of datasets. Summary of datasets grouped by imaging modality and stimulus type used or referenced in this work.

**Table 3.**
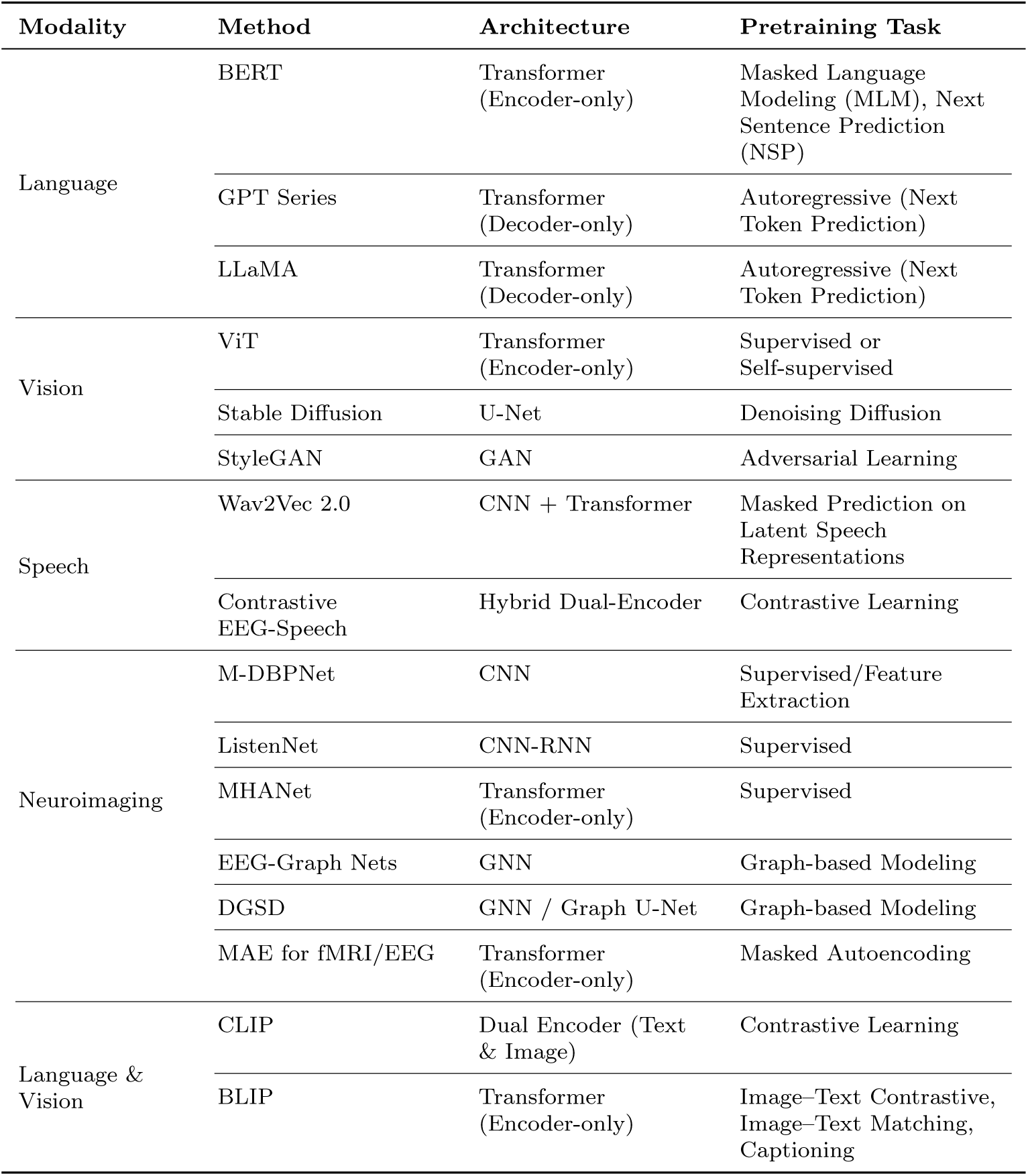
Overview of Key Foundation Model Architectures applied in BCI.

**Table 4.**
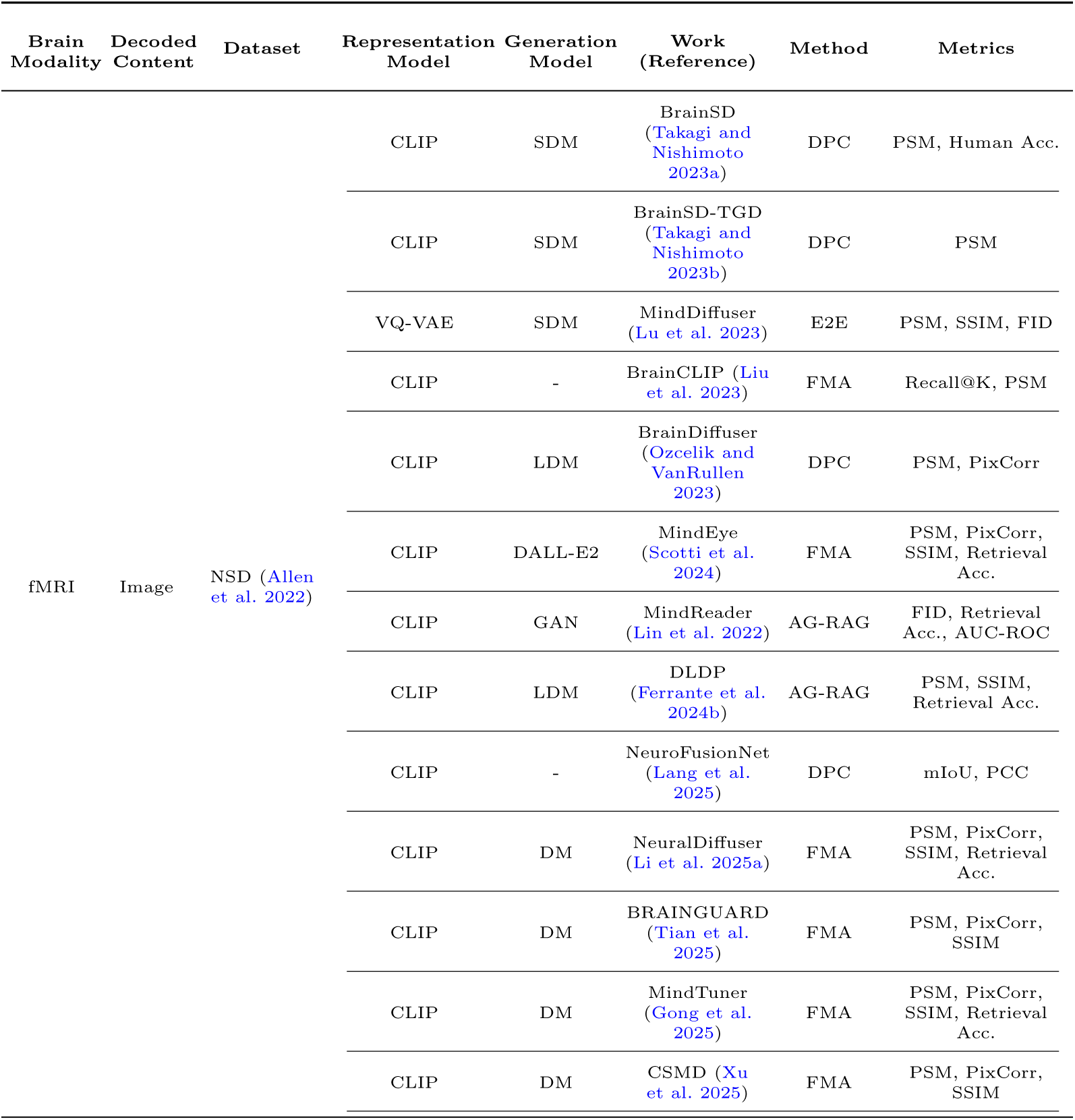
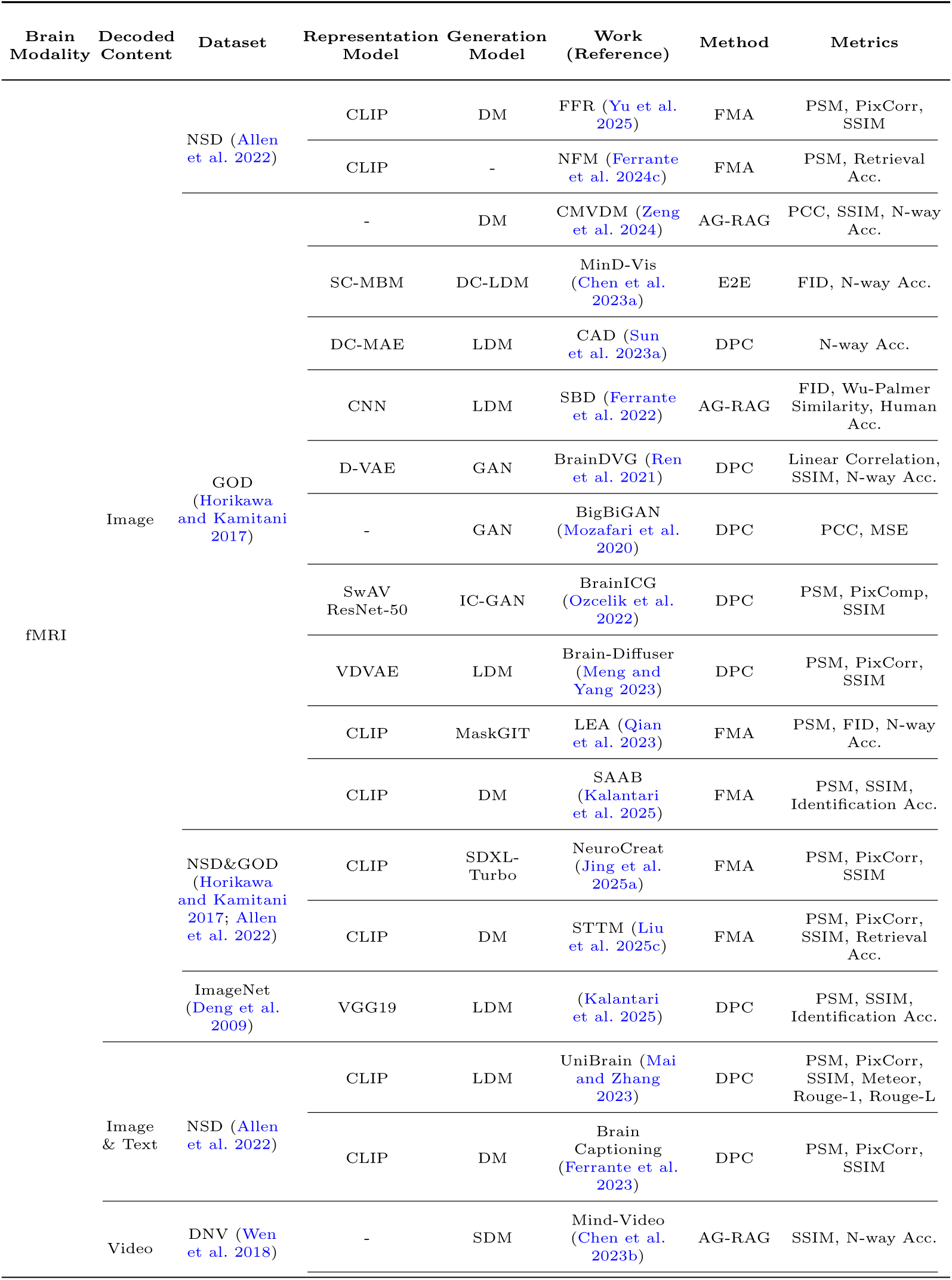
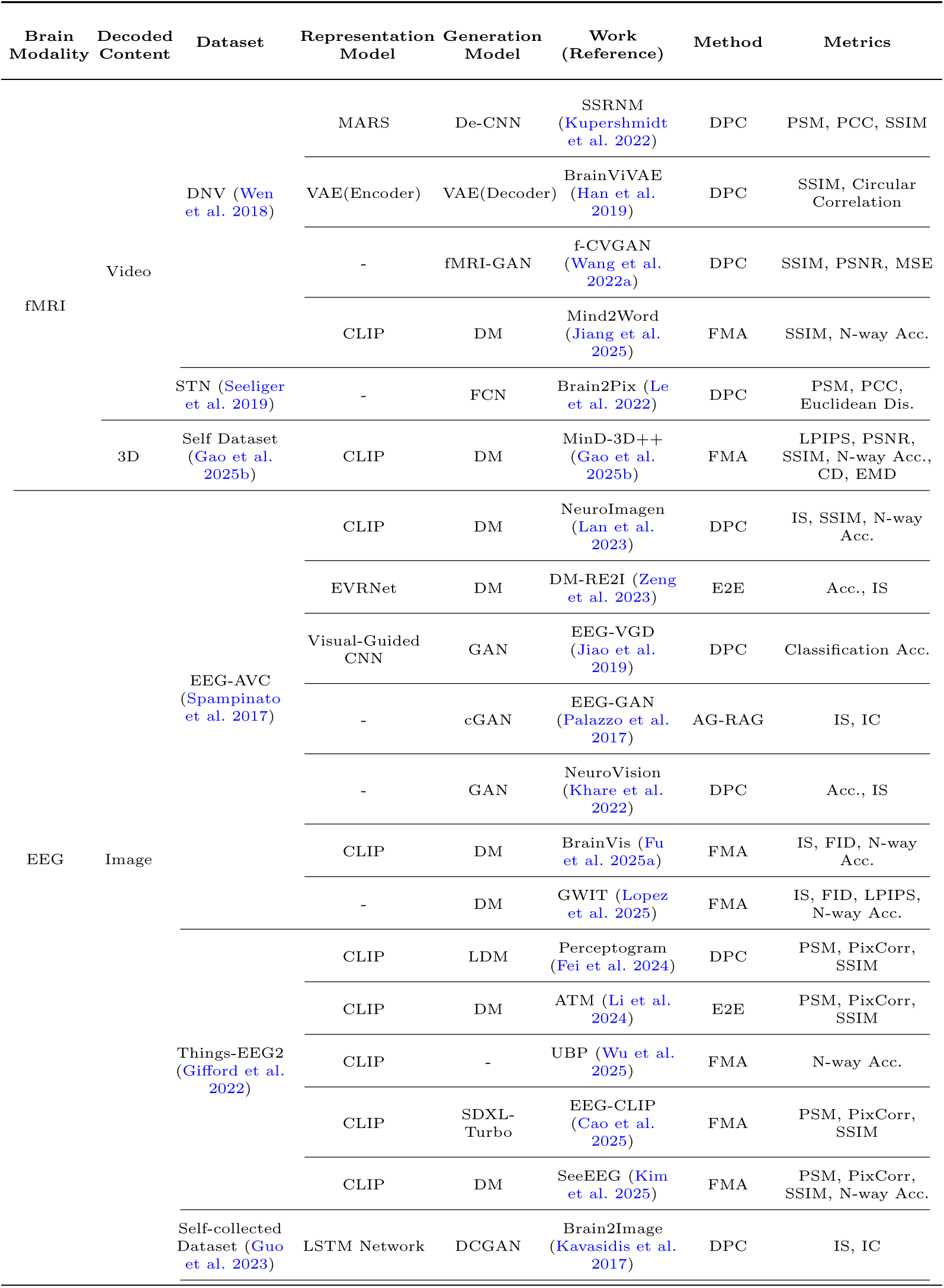
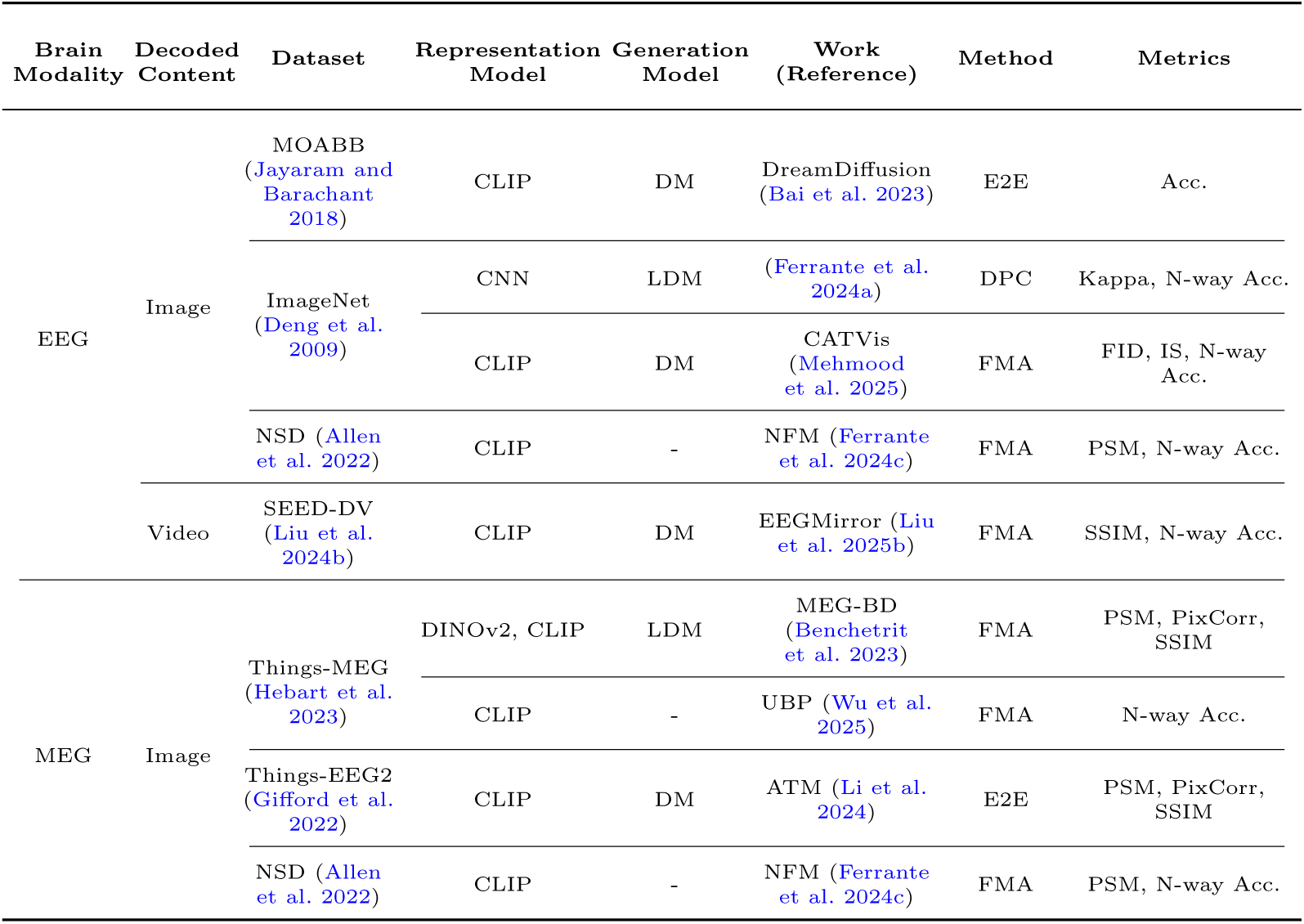
Summary of Decoding Visual Experiences (Images and Video). Columns denote: Brain Modality (neuroimaging method), Decoded Content (target output such as image or video), Dataset (benchmark used), Representation Model (feature extraction model), Generation Model (framework for reconstruction), Work (Reference), Method (Follows our taxonomy (DPC/FMA/E2E/AG-RAG/AG-EML)), Metrics (Perceptual Similarity Metrics (PSM) include CLIP, C3D (Convolutional 3D), AlexNet, Inception, EffNet-B, SwAV). FID is Fŕechet Inception Distance. mIoU is Mean Intersection over Union. PCC is Pearson Correlation Coefficient. CD is Chamfer Distance, EMD is Earth Mover’s Distance. IS is Inception Score. IC is Inception Classification Accuracy.

**Table 5.**
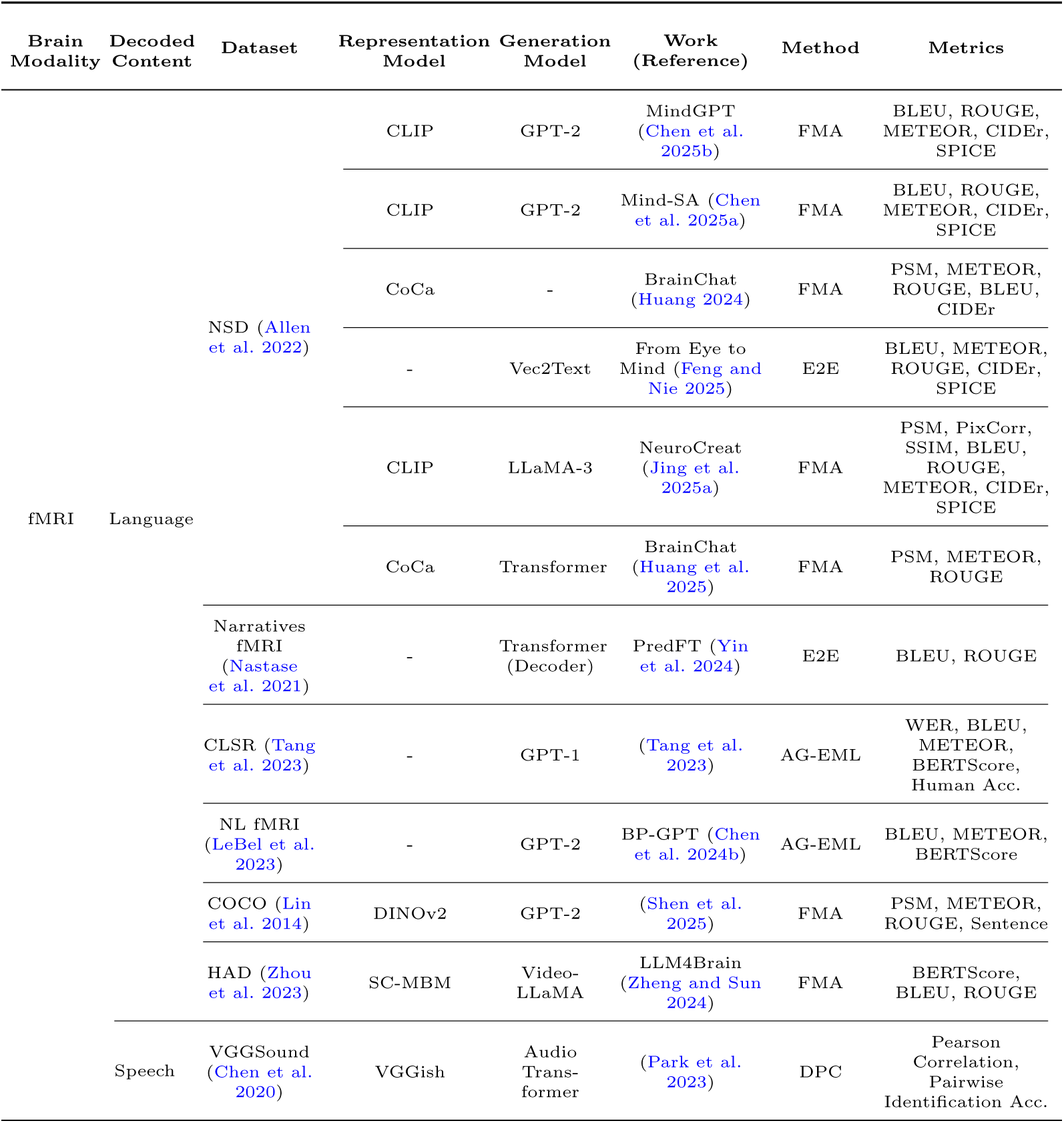
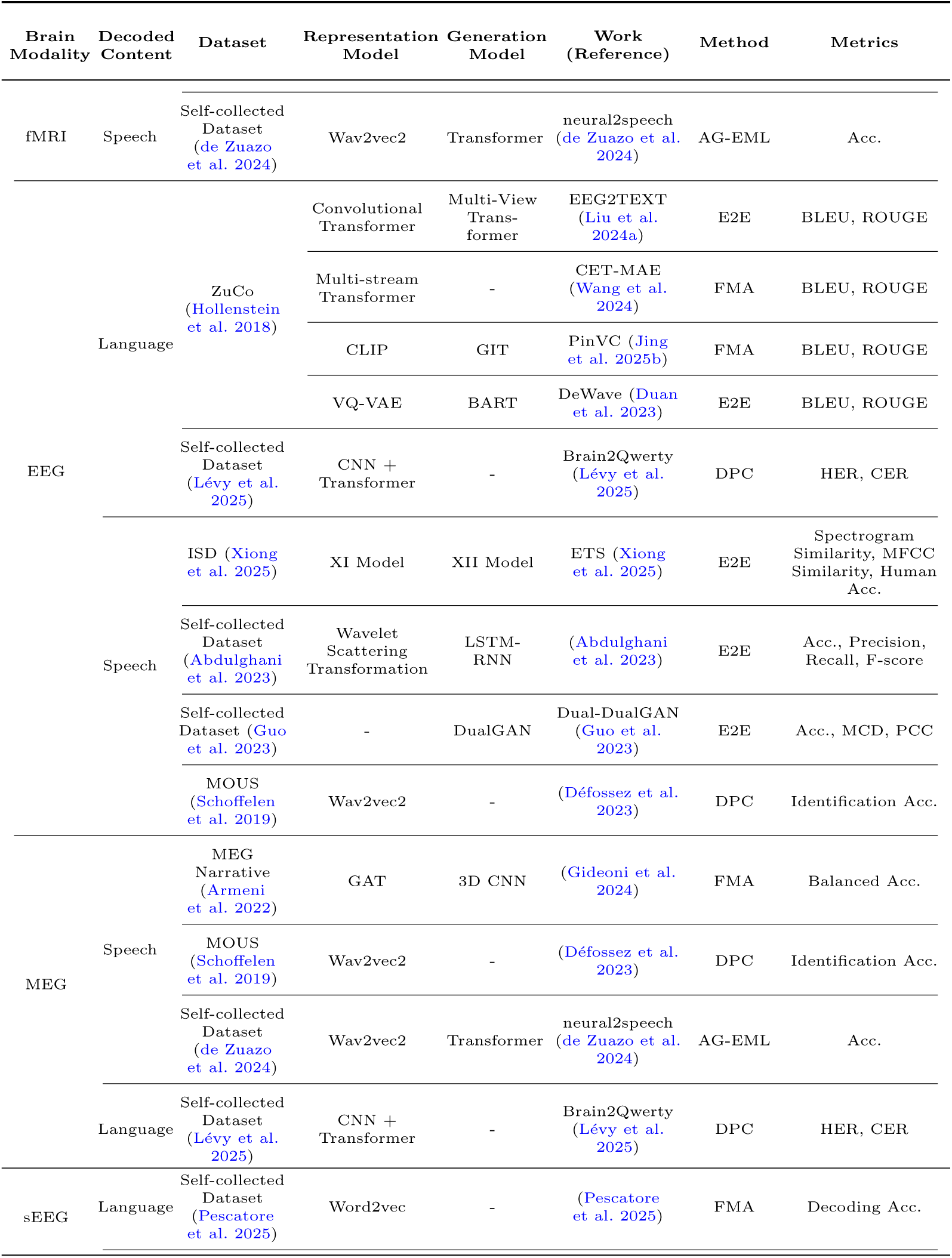
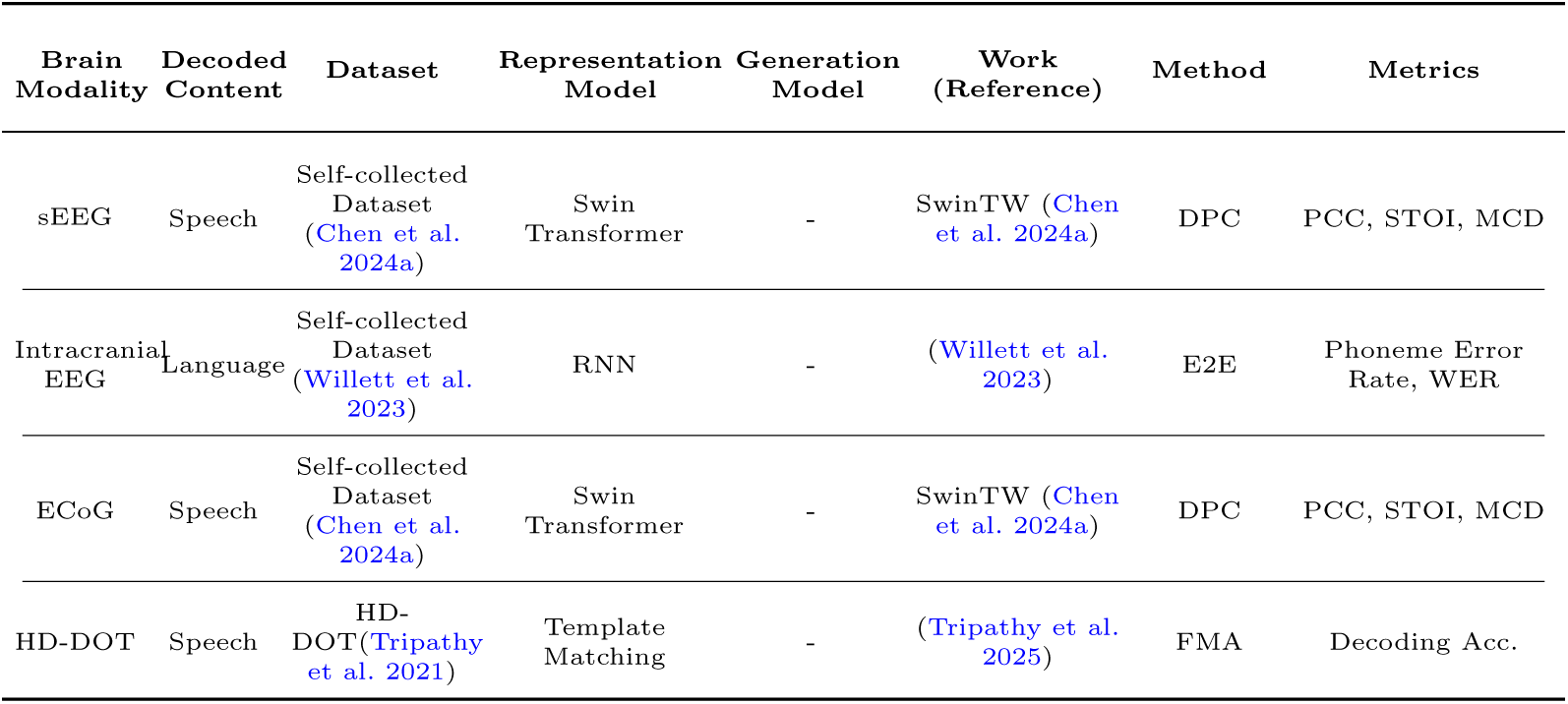
Summary Table of Recent Language/Speech Decoding Works. To help compare the upper limits of language/speech decoding, a small number of invasive studies were included as a reference, but these are not part of the core review scope of this table. Metrics(Perceptual Similarity Metrics (PSM) include CLIP, C3D (Convolutional 3D), AlexNet, Inception, EffNet-B, SwAV). WER is Word Error Rate. BLEU is Bilingual Evaluation Understudy. METEOR is Metric for Evaluation of Translation with Explicit ORdering. ROUGE is Recall-Oriented Understudy for Gisting Evaluation. CIDEr is Consensus-based Image Description Evaluation. SPICE is Semantic Propositional Image Caption Evaluation. MFCC is Mel Frequency Cepstrum Coefficients. MCD is Mel-cepstral distortion. PCC is Pearson Correlation Coefficient. HER is Hand Error Rate. CER is Character-error-rate. STOI is ShortTime Objective Intelligibility.

**Table 6.** Operating regimes and validation recommendations for FM-based and classical BCI pipelines. Acc: accuracy; ITR: information transfer rate; Corr: stimulus–response correlation; Fidelity metrics include SSIM/PSNR (vision) or WER/CER (speech); Semantic metrics include CLIPScore/BERTScore or retrieval-based consistency.

| Deployment Regime | Classical Baseline Strengths | FM-based Advantages | FM Failure Mode | Recommended Validation |
| --- | --- | --- | --- | --- |
| Low-latency closed-set control (P300/SSVEP/MI) | Stable Calibration; Low Compute | Marginal Benefit | Prompt/Decoding Sensitivity; Calibration Overhead | Cross-session / Acc-ITR / Latency |
| Small-data; High-noise; Online Drift | Sample-efficient; Robust Regularization | Priors help if aligned | Alignment Drift; Domain Shift | Cross-session / AUROC-Corr / Drift stress-test |
| Clinical Interpretability / Auditability | Transparent Assumptions; Inspectable Parameters | Semantic Scaffolding | Plausible but Unfaithful Generation | Cross-session / Fidelity+Semantic / Blinded human eval |
| Open-vocabulary Semantic Decoding (Naturalistic) | Limited Open-set Semantics | Strong Semantic Priors; Retrieval | Hallucination; Metric Mismatch | Cross-subject / Semantic+Fidelity / Prompt sensitivity |

**Table 7.** Prioritized research roadmap for foundation-model–based non-invasive brain decoding.

| Priority | Research Objective | Methodology | Reference Benchmarks |
| --- | --- | --- | --- |
| P1 | Benchmarking and comparability | Fixed splits; Matched baselines; Open evaluation scripts | NSD, THINGS, Narratives, Things-EEG2 |
| P2 | Cross-session and cross-subject robustness | Functional alignment; PEFT/adapters; Domain adaptation; Drift-robust training | Narratives, MOUS, HCP-style |
| P3 | Real-time and resource-constrained deployment | Streaming encoders; Distillation/quantization; Memory-efficient generation | AAD corpora, MOABB paradigms, portable EEG tasks |
| P4 | Trustworthy decoding and governance | Threat modeling; Red-teaming; Privacy-preserving learning | Any benchmark + Standardized misuse tests |

Throughout this survey, we distinguish controlled feasibility demonstrations from translation-level evidence. In particular, claims about broad generalizability are considered well supported only when results are reproduced under cross-session and cross-subject evaluation, and ideally under naturalistic/online constraints.

## 2 Theoretical Foundations: Bridging Computational Models and Biological Neural Processing

To understand how foundation models successfully decode brain activity, it is imperative to explore the underlying computational neuroscience theories. This section elucidates the mathematical and biological principles that justify the alignment between artificial architectures and neural substrates.

### 2.1 The Neural Coding Hypothesis and Decodability

The fundamental premise of brain decoding is the **Neural Coding Hypothesis**, which posits that mental states including perceptions, intentions, and semantic thoughts are not abstract entities but are represented through specific physical patterns of neural activity.

Mathematically, we define a mapping function *f* such that a stimulus or internal state **s**(*t*) yields a neural response **r**(*t*):

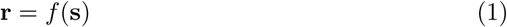

Decoding is feasible only if this mapping is systematic rather than stochastic. According to the Universal Approximation Theorem (Hornik et al. 1989), a neural network with sufficient capacity can approximate any continuous function. Thus, if the encoding process *f* is information-preserving, there exists an inverse mapping *g* such that:

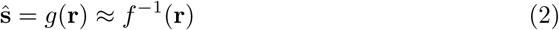

Beyond representational capacity, the Statistical Identifiability of the neural signal is crucial. For distinct mental states **s**_1_ *̸*= **s**_2_, the conditional distributions must satisfy *p*(**r**|**s**_1_) *̸*= *p*(**r**|**s**_2_). This discriminability forms the basis of Bayesian optimal decoding, where the model seeks to approximate the posterior arg max**_s_** *p*(**s**|**r**) (Paninski et al. 2007).

### 2.2 Representational Geometry and Isomorphism

Recent success in utilizing Large Language Models (LLMs) for decoding stems from the theory of Geometric Isomorphism. This theory suggests that the relational structure of neural representations mirrors the structure of semantic embeddings:

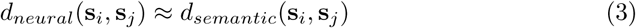

When the distance metrics between neural manifolds and artificial latent spaces are preserved, simple linear transformations (e.g., ridge regression) suffice to align the two domains (Kriegeskorte et al. 2008). This isomorphism explains why high-dimensional semantic embeddings from foundation models can serve as effective proxies for biological neural representations (Haxby et al. 2001).

### 2.3 Key Theoretical Perspectives on Neural Processing

We categorize the current computational understanding of brain-model alignment into three primary schools of thought:

- **Bayesian and Predictive Coding:** Proponents of this view, such as (Friston 2009) and (Bottemanne 2025), argue that the brain functions as a hierarchical generative model. It minimizes ”surprisal” by constantly updating internal beliefs through probability inference. In this context, decoding is the process of extracting the brain’s internal ”prior” and ”prediction error” signals, which can be mapped to synaptic plasticity and Bayesian updates.
- **Traditional Neural Coding:** This school focuses on the structural and temporal properties of neuronal population activity. Works by (Olshausen and Field 2004) emphasize *sparse coding*, while (Tsien 2016) introduces the ”Power-of-N” combinatorial logic. Notably, Tsien suggests that ”rare silence” (synchronized inactive states) within neuronal cliques may carry primary information, providing a new perspective on neural variability and self-information encoding that transcends simple rate coding.
- **Goal-Driven Artificial Neural Networks (ANNs):** This perspective, championed by (Yamins and DiCarlo 2016) and (Eickenberg et al. 2017), treats deep networks as functional surrogates for the brain. It posits that optimizing a model on a complex task (e.g., object recognition or next-token prediction) naturally leads to internal representations that are hierarchical and functionally similar to those found in the human cortex, even if the biological implementation (spikes vs. gradients) differs.

## 3 The Landscape of Non-Invasive Brain Decoding: Modalities and Challenges

To appreciate the potential impact of foundation models, it is essential to first understand the specific characteristics of non-invasive brain decoding techniques and the fundamental challenges they present (Summarized in Table 1).

### 3.1 Non-Invasive Neuroimaging Modalities

Non-invasive recordings in BCI and brain decoding are dominated by EEG, MEG, fMRI, and fNIRS, which differ in their physical principles and therefore in temporal/spatial resolution, robustness to noise and motion, cost, and deployability. These differences directly shape what can be decoded, how models should be designed, and how results are ultimately interpreted.

**Electroencephalography (EEG)** captures voltage fluctuations on the scalp generated by synchronous activity of large neuronal populations (da Silva 2013; Buzśaki et al. 2012). Its chief advantage is millisecond-level temporal resolution, which enables tracking rapid neural dynamics and supports closed-loop or real-time applications (Zhu et al. 2024). EEG systems are relatively inexpensive, portable, and easy to deploy, contributing to their ubiquity in research and clinical settings. At the same time, volume conduction blurs spatial specificity, typically limiting spatial resolution to the centimeter scale (da Silva 2013); recordings are also vulnerable to physiological artifacts (eye blinks, muscle activity) and environmental electrical noise (Zhang et al. 2021).

**Magnetoencephalography (MEG)** measures the minute magnetic fields produced by the same neural currents that underlie EEG signals (da Silva 2013; Buzśaki et al. 2012). Because magnetic fields are less distorted by skull and scalp than electric potentials, MEG generally offers better source localization (millimeters to centimeters) while preserving millisecond temporal precision (da Silva 2013). These benefits come with practical constraints: MEG systems are expensive, require magnetically shielded environments, and are sensitive to head motion, which limits portability and broad accessibility.

In contrast to the electrophysiological modalities, **Functional Magnetic Resonance Imaging (fMRI)** infers neural activity indirectly via blood-oxygen-level-dependent (BOLD) signals (Logothetis 2008). The technique excels in spatial resolution at the millimeter scale, enabling precise mapping of cortical and subcortical representations and supporting decoding of fine-grained spatial patterns such as visual feature maps (Logothetis 2008; Naselaris et al. 2009; Kay et al. 2008). Its principal limitation is temporal: the hemodynamic response peaks several seconds after neural events, resulting in second-level temporal resolution (Logothetis 2008). Practical considerations—high scanner cost, immobility, acoustic noise, and strict motion constraints—further narrow its real-time applicability.

**Functional Near-Infrared Spectroscopy (fNIRS)**, which estimates changes in oxy- and deoxy-hemoglobin using near-infrared light, shares the vascular origin of fMRI signals while offering greater portability and lower cost (Ferrari and Quaresima 2012). It provides an intermediate operating point with temporal resolution on the order of seconds and spatial resolution at the centimeter scale, and it is relatively tolerant to modest movement compared with EEG or fMRI (Ferrari and Quaresima 2012). However, limited penetration depth restricts sensitivity primarily to superficial cortical tissue, and spatial precision and coverage remain below those of fMRI (Ferrari and Quaresima 2012).

Taken together, these modalities impose fundamental trade-offs (Table 1). High temporal resolution in EEG and MEG is advantageous for rapid interaction and control, but weaker spatial localization complicates decoding of fine spatial structure. fMRI provides the inverse profile—excellent spatial precision that supports detailed representational decoding, offset by poor temporal resolution that hinders fast feedback. fNIRS occupies a pragmatic middle ground, improving deployability while rarely achieving the best-in-class performance in either domain (Ferrari and Quaresima 2012). These properties dictate modeling priorities: EEG/MEG pipelines benefit from sophisticated temporal processing and artifact mitigation, whereas fMRI pipelines emphasize high-dimensional spatial feature extraction and alignment. Ultimately, the ceiling on any decoding approach—including those leveraging foundation models—is tightly coupled to the information content and quality of the chosen signal source (da Silva 2013; Logothetis 2008; Ferrari and Quaresima 2012). Figure 1 summarizes the typical workflow for non-invasive brain data acquisition and preprocessing using fMRI, EEG, and MEG. This figure not only illustrates how raw measurements are transformed into analyzable time series and brain maps (such as BOLD time series, ERP/ERF), but also highlights the core preprocessing steps for different modalities. These processed outputs serve as direct inputs for downstream FM-based decoders during training and inference, thus supporting the discussion of the tradeoffs in Table 1 at the data level.

### 3.2 Fundamental Challenges in Non-Invasive Decoding

Beyond modality-specific limitations, non-invasive brain decoding faces a set of crosscutting obstacles that fundamentally constrain robustness and scalability (Abiri et al. 2019; Luo et al. 2024; Geirnaert et al. 2021; Shirakawa et al. 2024). First, signals are weak and easily corrupted: propagation through tissue attenuates and distorts neural activity, while eye blinks, muscle activity, and environmental interference introduce non-neural components that are difficult to remove without damaging informative structure (Zhang et al. 2021; Geirnaert et al. 2021). Second, the data are high-dimensional, non-stationary, and idiosyncratic—varying across and within individuals due to anatomy, physiology (fatigue, arousal), cognitive state (attention), and task demands—so models that perform well in one session or subject often fail to generalize without costly recalibration (Luo et al. 2024; Chen et al. 2023a). Third, hard physical limits in spatial and temporal resolution cap extractable information: slow hemodynamics blur rapid events in fMRI (Logothetis 2008), while volume conduction in EEG mixes sources and degrades spatial separability (da Silva 2013).

Practical deployment adds further constraints. Many applications require end-to-end latencies on the order of hundreds of milliseconds, forcing a difficult accuracy, latency, compute trade-off, particularly on portable or embedded hardware (Zink et al. 2017). Ecological validity is another barrier: models tuned on tightly controlled laboratory paradigms (e.g., P300 (Pritchard 1981; Solon et al. 2019), SSVEP (Zhu et al. 2010), or motor imagery (Altaheri et al. 2023; McGeady et al. 2019)) often degrade in naturalistic settings with unconstrained behavior, rich sensory input, and movement, creating a persistent “lab-to-life” gap (Luo et al. 2024; Abiri et al. 2019). Finally, data scarcity remains chronic: collecting large-scale, well-annotated non-invasive datasets is expensive and time-consuming, limiting the training of capacity-hungry models and amplifying risks of spurious correlations and distribution shift (Jiang et al. 2024; Shirakawa et al. 2024).

These challenges jointly shape algorithmic priorities for non-invasive decoding: aggressive but principled denoising and artifact suppression; adaptation to non-stationarity and cross-subject variability; architectures that respect each modality’s information bottlenecks; streaming inference with bounded latency; and training regimes that mitigate data scarcity (e.g., self-supervision, cross-subject/domain adaptation, and careful regularization). The attainable performance of foundation-model-based approaches is therefore tightly coupled to how effectively they confront these constraints.

While we have discussed accidental artifacts such as those caused by ocular, muscular, or environmental interference, the real-world deployment of BCI technology also introduces the potential for intentional corruption. Recent studies highlight adversarial perturbations; small, structured injections of noise which are optimized to mislead a trained model and induce large decoding errors (Zhang et al. 2020; Lubars and Chin 2022; Li et al. 2021). As the field shifts towards large foundation models, these threats to data integrity extend from test-time evasion to backdoor attacks, where triggers are implanted during pre-training, enabling specific mis-decoding to occur when a trigger pattern, such as an adversarial perturbation, is present (Meng et al. 2022). This distinction matters because robustness to incidental artifacts/’noise’ does not imply robustness to intentional perturbations, which have been shown to bypass standard denoising pipelines remaining undetected (S-a et al. 2020).

### 3.3 Limitations of Pre-Foundation Model Approaches

Traditional computational methods often struggled to fully overcome these challenges. Early approaches using linear models (e.g., LDA, linear SVM, regression) (De Taillez et al. 2020; Wong et al. 2018) were efficient but lacked capacity for complex non-linear neural dynamics. For instance, linear models in Auditory Attention Decoding (AAD) (e.g., SR, TRF, CCA) (Wong et al. 2018; Fiedler et al. 2017; De Cheveigńe et al. 2018; Dmochowski et al. 2018) showed declining performance in short time windows crucial for real-time use (Cai et al. 2020; Vandecappelle et al. 2021).

Deep learning (CNNs, RNNs) offered improvements by automatically learning features and temporal dependencies (Craik et al. 2019; Zhou et al. 2018), enhancing tasks like motor imagery classification (Altaheri et al. 2023), ERP detection (Solon et al. 2019), fine-grained semantic decoding (Wang et al. 2020), and even early visual reconstruction (Nishimoto et al. 2011; Beliy et al. 2019; Shen et al. 2019). However, these models, often trained from scratch on limited BCI datasets, still struggled with generalization due to data scarcity and variability (Luo et al. 2024), had difficulty decoding complex semantic content and often required extensive preprocessing (Craik et al. 2019).

### 3.4 The Promise of Foundation Models for BCI Challenges

Foundation models (FMs) offer a principled route to address several long-standing hurdles in non-invasive decoding by leveraging large-scale pretraining and transfer learning. Trained on vast neuroimaging or multimodal corpora, FMs can internalize the statistical structure of brain signals and their relations to language and vision, thereby providing robust priors for downstream decoding under data and noise constraints.

First benefit concerns low SNR and artifacts. Self-supervised objectives—such as masked reconstruction in Masked Autoencoders (MAEs)—encourage models to capture invariant structure in unlabeled recordings, improving denoising and feature extraction without relying on brittle preprocessing (Zhang et al. 2021; He et al. 2022; Chen et al. 2023a). Because these objectives learn to predict missing or corrupted segments from context, they naturally attenuate physiological and environmental noise while preserving informative components of the signal.

Second, scale and diversity in pretraining mitigate signal complexity and variability across subjects, sessions, and tasks. By exposing encoders to heterogeneous distributions, FMs learn more transferable representations that reduce subject-specific calibration and improve cross-dataset generalization; fine-tuning or lightweight adaptation then specializes the model to new cohorts with limited labels (Luo et al. 2024; Tang et al. 2023). This approach directly targets the non-stationarity that undermines conventional pipelines.

Third, although FMs cannot break the physical limits of spatial or temporal resolution, their capacity supports richer integration across sensors, space, and time—amortizing information over extended contexts to infer latent neural states that are only weakly expressed in raw measurements. In parallel, pretrained semantic spaces from language and vision (e.g., LLMs, CLIP) provide high-level targets that align with cortical representations (Radford et al. 2021; Mitchell et al. 2008; Huth et al. 2016; Gong et al. 2024). Mapping brain activity into these structured manifolds enables more expressive decoding—such as detailed image reconstructions or continuous speech content—than task-specific classifiers alone (Scotti et al. 2024; Liu et al. 2023; Tang et al. 2023). When coupled with conditional generators (diffusion models, autoregressive LLMs), strong priors can produce plausible outputs from noisy inputs, effectively regularizing the ill-posed inverse problem (Takagi and Nishimoto 2023a; Radford et al. 2019; Gong et al. 2024).

Finally, FMs help ameliorate data scarcity. Large-scale pretraining on web-scale text/images and emerging brain datasets bootstraps powerful initializations that require far fewer task-specific labels to reach competitive performance, and self-supervised learning on unlabeled neurodata further expands usable training signal (Jiang et al. 2024; Gong et al. 2024; Jayalath et al. 2024). In essence, by importing knowledge from broader corpora and aligning it with neural measurements, FMs offer a unifying recipe—robust representation learning, transferable adaptation, semantic alignment, and generative priors—that can accelerate progress toward reliable, non-invasive BCI systems.

### 3.5 Datasets & Benchmarks

In non-invasive brain decoding research, commonly used datasets can be systematically divided according to stimulus type, including image, video, speech/audio, reading/language, and multi-modal types. Different types of stimuli correspond to different neural representation characteristics and experimental paradigms, covering multiple levels from low-level vision to high-level semantic processing. Table 2 provides an overview of the datasets used in this article, along with their key elements (Dataset, Modality, Task/Stimuli, Standard BCI Paradigm, Research Paradigm, Subjects), to facilitate quick location and comparison.

#### 3.5.1 Image Stimulus

Image datasets are fundamental to visual decoding research, primarily focusing on static scene and object representations. Typical datasets include NSD (Allen et al. 2022) and GOD (Horikawa and Kamitani 2017), which are used for high-resolution brain imaging of natural scenes and category recognition, respectively; EEG-AVC (Spampinato et al. 2017) and Things-EEG2 (Gifford et al. 2022) utilize high-temporal resolution EEG to record rapid visual categorization. Things-MEG (Hebart et al. 2023) extends the same task to MEG, providing millisecond-level temporal dynamics. These datasets often employ the paradigm of subjects viewing natural or object images and are suitable for training and validating visual encoding and decoding models.

#### 3.5.2 Video Stimulus

Video datasets emphasize temporal information and ecological validity. These include DNV (Wen et al. 2018), STN (Seeliger et al. 2019), and HAD (Zhou et al. 2023) in fMRI, which are used for natural scene or action recognition tasks. In EEG, SEED-DV (Liu et al. 2024b) combines rich semantic annotations with multimodal signals for semantic decoding from EEG to video. MEG Narrative (Armeni et al. 2022) and HD-DOT (Tripathy et al. 2021) provide natural viewing data using high temporal resolution and high-density optical imaging. This type of video data supports temporal modeling, event boundary recognition, and representation learning of complex visual semantics.

#### 3.5.3 Audio Stimulus

Auditory and speech datasets typically focus on natural narratives. For example, CLSR (Tang et al. 2023), NL fMRI (LeBel et al. 2023), and Narratives fMRI (Nastase et al. 2021) use fMRI to record brain activity while participants listen to natural stories. The Narratives fMRI dataset (Nastase et al. 2021) is large (345 subjects), making it a landmark resource for natural language decoding. The EEG Imagined Speech Dataset (ISD) (Xiong et al. 2025) covers both real and imagined speech tasks, supporting end-to-end research on direct speech generation from brain signals.

#### 3.5.4 Reading/Language Stimulus

Reading and language datasets focus on the neural mechanisms of language processing. ZuCo (Hollenstein et al. 2018) combines EEG and eye tracking to record brain activity while participants read natural sentences. MOUS (Schoffelen et al. 2019) and Narratives fMRI (Nastase et al. 2021) cover reading and auditory language tasks, and provide structural and diffusion MRI for multimodal fusion. ISD (Xiong et al. 2025) involves speech imagery and language production, and is suitable for brain decoding tasks related to language production.

#### 3.5.5 Multi-stimuli

Some datasets cover a variety of stimulus and task types, supporting multi-paradigm comparisons and cross-modal studies. For example, both HCP (Elam et al. 2021) and MOUS (Schoffelen et al. 2019) offer large-scale structural and functional MRI (HCP includes 1,200 subjects), covering multiple tasks such as reading, hearing, and vision, and providing additional information such as genetics and behavioral data. MOABB (Jayaram and Barachant 2018) integrates multiple public EEG datasets and standardizes paradigms such as MI, ERP, and SSVEP, making it a valuable resource for algorithm comparison and benchmarking.

## 4 Foundation Models: Core Concepts and Architectures for Brain Decoding

Understanding the fundamental principles and specific architectures of FMs is crucial for appreciating their application in non-invasive brain decoding.

### 4.1 Defining Foundation Models

Foundation Models (FMs) refer to a class of large-scale artificial intelligence models that are trained on massive and diverse datasets—often using self-supervised learning objectives—and subsequently adapted to a wide range of downstream tasks through fine-tuning or prompting strategies (Radford et al. 2021; Devlin et al. 2019). Unlike traditional task-specific models, FMs acquire general-purpose representations that can be reused across domains and modalities, making them particularly powerful in settings where labeled data is scarce or heterogeneity is high.

At their core, FMs combine scale with broad pre-training. Parameter counts span from hundreds of millions to trillions, and training data frequently reach web or multi-modal scale; such regimes enable models to internalize high-order statistical regularities and support emergent capabilities beyond those of smaller counterparts (Kaplan et al. 2020; Henighan et al. 2020; Hestness et al. 2017). Pre-training objectives instantiate this breadth: masked-token prediction in language (e.g., BERT) and cross-modal contrastive alignment in vision–language (e.g., CLIP) endow models with semantic structure, syntactic competence, and cross-modal correspondences without manual annotation (Devlin et al. 2019; Radford et al. 2021).

Equally central is adaptability. Because pre-training yields versatile representations, modest amounts of task-specific data often suffice to specialize an FM for classification, generation, retrieval, or reasoning under few-/zero-shot settings. In neuroimaging, this adaptability directly addresses chronic data scarcity and subject variability by enabling lightweight transfer to new cohorts, sessions, or paradigms (Tang et al. 2023).

Prominent exemplars include large language models (GPT (Radford et al. 2019), BERT (Devlin et al. 2019)), vision–language encoders (CLIP (Radford et al. 2021)), and modern generative models (e.g., diffusion backbones used for image synthesis and reconstruction (Takagi and Nishimoto 2023a)). Conceptually, the paradigm shifts from crafting bespoke pipelines to leveraging a powerful, pre-trained base that can be aligned with specific BCI objectives through fine-tuning or prompting.

### 4.2 Key Properties Driving BCI Advancement

Foundation models advance non-invasive decoding chiefly through three mutually reinforcing properties. First, they learn high-quality, hierarchical representations that align with neural population codes. Vision and language FMs capture semantic structure across layers (Radford et al. 2021, 2019); these internal embeddings correlate with brain activity during corresponding tasks—e.g., fMRI responses in visual cortex track layers of vision encoders and vision–language models such as CLIP, whose embedding space aligns with neural responses to images and text (Mitchell et al. 2008; Radford et al. 2021; Liu et al. 2023; Gong et al. 2024). Beyond holistic embeddings, recent work demonstrates that disentangling conceptual representations into fine-grained semantic subdimensions can significantly enhance the interpretability and fidelity of mapping neural activity to semantic meaning (Zhang et al. 2025). Notably, larger language models often exhibit stronger brain alignment, suggesting scaling benefits for representational fidelity (Antonello et al. 2023; Gao et al. 2025a). In addition to unsupervised language models based on English, research on Chinese has also shown that efficient parameter tuning methods can significantly affect language-brain alignment (Zhang et al. 2025).

Second, broad pre-training underwrites transfer and generalization. Because FMs are exposed to heterogeneous, large-scale corpora, their features transfer with modest adaptation to small BCI datasets, improving robustness across subjects, sessions, and paradigms and mitigating the chronic data-scarcity problem (Luo et al. 2024; Tang et al. 2023; Li et al. 2024). This adaptability enables few-/zero-shot behaviors that are difficult to obtain with narrowly trained decoders.

Third, strong conditional generation closes the loop from brain signals to rich outputs. Diffusion (Takagi and Nishimoto 2023a) backbones and autoregressive LLMs (Radford et al. 2019) can be conditioned on FM-derived neural embeddings to synthesize images or text, using powerful priors to regularize the ill-posed inverse problem and produce plausible reconstructions even under noisy conditioning (Chen et al. 2023b; Sun et al. 2023a; Chen et al. 2023a; Scotti et al. 2024). Together, neural alignment, transferable representations, and generative priors create an end-to-end pathway from weak non-invasive signals to high-level percepts and concepts, thereby accelerating progress in reliable BCI decoding (Gong et al. 2024).

### 4.3 Relevant Architectures and Their Roles in BCI

Neuroimaging-native architectures exploit the structure of brain data itself. Purpose-built encoders (e.g., M-DBPNet, ListenNet, MHANet) tailor feature extraction to fMRI/EEG characteristics, while graph-based models (e.g., EEG graph networks, DGSD) encode inter-sensor topology and functional connectivity to better capture mesoscale dynamics (Fan et al. 2025b,a; Li et al. 2025b; Fan et al. 2024). Together, these modality-specific roles illustrate how FMs function not as a monolith but as a toolbox: language decoders contribute generative priors, vision encoders and contrastive bridges supply structured semantic targets, speech encoders provide acoustically grounded manifolds, and neuroimaging models respect the physics and topology of the measurements—collectively advancing the fidelity and robustness of non-invasive decoding.

Recent advances in non-invasive brain decoding have been propelled by the targeted adoption of diverse foundation-model (FM) architectures across language, vision, speech, and neuroimaging, each filling a distinct role rather than occupying the same rung in a uniform taxonomy (Table 3). In language, transformer families split naturally into encoder-only and decoder-only uses. Encoder-only models such as BERT provide context-rich embeddings via masked-token objectives that serve as stable targets or intermediate representations for textual decoding and semantic alignment (Devlin et al. 2019). Decoder-only models—including GPT and LLaMA—offer strong autoregressive priors that support coherent sequence generation from neural features, enabling reconstruction or continuation of linguistic content under noisy or partial conditioning (Radford et al. 2019; Touvron et al. 2023).

In vision, encoder-centric ViTs and masked-autoencoding regimes (e.g., MAE) learn robust patch-level structure that is well matched to spatially distributed neural signals, especially for fMRI-based decoding where fine-grained representational geometry matters (Dosovitskiy et al. 2020; He et al. 2022). Dual-encoder models such as CLIP align images and text through contrastive learning and thus provide a shared semantic manifold that links brain activity to multimodal concepts, improving both zero-/few-shot decoding and retrieval (Radford et al. 2021). For generative targets, diffusion backbones now dominate high-fidelity reconstructions from brain activity, superseding earlier GAN-based (Goodfellow et al. 2020) approaches for naturalistic images by leveraging powerful priors and iterative denoising.

For speech, self-supervised audio encoders like Wav2Vec 2.0 capture latent phonetic and prosodic structure through masked prediction on large unlabeled corpora, yielding representations that can be aligned to EEG/MEG correlates of auditory perception or imagery (Baevski et al. 2020). Complementary contrastive dual-encoder formulations map speech and neural activity into a shared space, facilitating recognition or retrieval of perceived/imagined speech with minimal labels (Qiu et al. 2025b).

## 5 Foundation Models Driving Advancements in Non-Invasive Brain Decoding

The integration of FMs has catalyzed significant progress, achieving unprecedented accuracy and richness in interpreting brain signals related to vision, language, speech, and auditory attention. A common conceptual pipeline involves: 1) Extracting denoised representations from neural signals (often using MAEs or SSL (Zhang et al. 2021; Chen et al. 2023a; Jiang et al. 2024; Jayalath et al. 2024)), 2) Connecting neural signals with decoding targets by aligning neural representations with FM semantic spaces (e.g., CLIP, LLM embeddings) using mapping modules and contrastive learning (Liu et al. 2023; Tang et al. 2023; Li et al. 2024; Chen et al. 2024b), and 3) Generating decoding results using conditional FMs (Diffusion Models, LLMs) guided by the aligned neural representations, leveraging strong priors for high-quality output (Chen et al. 2023b; Sun et al. 2023a; Chen et al. 2023a; Scotti et al. 2024). To provide a context for recent research, Figure 2 maps representative works from 2017–2025 across modalities and output types; we use it as a roadmap for the subsections that follow.

### 5.1 Foundation Models in Neuroimaging

In non-invasive brain decoding, foundation models (FMs) typically play three complementary roles: First, they utilize self-supervised/weakly supervised methods to obtain transferable representations, such as MAE (He et al. 2022), wav2vec 2.0 (Baevski et al. 2020), and BLIP/CLIP (Li et al. 2022), providing a robust foundation for small sample sizes and cross-subject learning (Radford et al. 2021; Rombach et al. 2022). Second, they map fMRI/EEG/MEG to the semantic or acoustic manifolds of pre-trained models through neuro-semantic/acoustic alignment. For example, aligning brain representations to the visual-linguistic space of CLIP/BLIP or the acoustic space of wav2vec 2.0 (Liu et al. 2023; Radford et al. 2021; Rombach et al. 2022). Third, they leverage strong generative priors (diffusion/autoregression) to transform weak and time-delayed neural conditions into readable image, text, and speech outputs (Rombach et al. 2022; Takagi and Nishimoto 2023a; Ozcelik and VanRullen 2023; Lu et al. 2023). A representative approach in recent years is ”fMRI → CLIP semantic embedding → latent space diffusion reconstruction,” which significantly outperforms earlier task-specific networks in terms of semantic consistency and perceptual quality (Tang et al. 2023; Liu et al. 2023). Meanwhile, native neuroimaging models (such as M-DBPNet (Fan et al. 2025b), ListenNet (Fan et al. 2025a), MHANet (Li et al. 2025b), EEG graph networks/self-distilled graph networks, and MAE for fMRI/EEG) explicitly encode sensor topology and measurement physics to compensate for the modal gap in general FMs. Overall, the triadic collaboration of transferable representation + neural-semantic alignment + generative priors is connecting non-invasive weak signals end-to-end to high-level readable content, demonstrating stable gains in small-sample, cross-subject, and zero/small-sample scenarios.

At the same time, rapidly evolving role for foundation models focuses on learning universal representations of neuro-neuronal connectivity. These models, often termed Neuro-Connectome Transformers or Graph Foundation Models, directly consume fMRI/EEG time-series or derived functional connectivity matrices. Instead of learning semantic relationships like LLMs, their primary objective is to develop generalized feature extractors that capture the complex, dynamic linking mechanisms and rules between distinct neural populations (Shehzad et al. 2024; Kim et al. 2023). This approach often leverages Graph Transformer architectures to model the brain as a dynamic graph, employing self-attention mechanisms to discover high-order, global dependencies across brain regions, leading to powerful, transferable representations for tasks like brain disorder diagnosis and biotyping, even across large-scale heterogeneous datasets (Wei et al. 2025). Furthermore, techniques like Masked Brain Modeling (MBM), applied directly to raw fMRI data, serve as a potent form of self-supervised learning to force the model to infer and thus implicitly encode the underlying functional relationships between masked and unmasked brain areas (Chen et al. 2023a). This paradigm shift moves neuroimaging analysis from fixed, statistical connectivity measures towards flexible, learnable, and generalizable neuro-relational embeddings.

### 5.2 Decoding Visual Experiences

Visual decoding seeks to reconstruct perceived images or videos from brain activity by coupling neural embeddings with powerful priors from foundation models. Between 2023 and 2025, progress has accelerated as CLIP-style semantic spaces and diffusion backbones became standard components across modalities (overview in Table 4; schematic in Figure 3). In fMRI, spatial resolution is key (Liu et al. 2025a). The dominant strategy combines CLIP’s representational power with diffusion models’ generative prowess (Gong et al. 2024). An encoder maps fMRI signals to CLIP’s shared embedding space, capturing semantics (Gong et al. 2024; Liu et al. 2023). These embeddings condition a latent diffusion model (e.g., Stable Diffusion (Takagi and Nishimoto 2023a)) to generate the image (Scotti et al. 2024; Gong et al. 2024; Sun et al. 2023b). CLIP provides semantic guidance, while diffusion models ensure fidelity (Chen et al. 2023a; Gong et al. 2024). Foundational work like MinD-Vis (CVPR 2023) used sparse masked modeling (MAE-like) for fMRI representation and a double-conditioned LDM (DC-LDM) for reconstruction, addressing variability (Chen et al. 2023a). Brain-Diffuser predicted multimodal CLIP features from fMRI to condition Versatile Diffusion. These FM-based methods yield significant improvements in semantic accuracy and perceptual fidelity over earlier approaches (Chen et al. 2023a; Liu et al. 2025a).

With EEG, the emphasis shifts to exploiting millisecond dynamics under limited spatial precision. Recent end-to-end schemes learn a projection from EEG to CLIP space—e.g., via an Adaptive Thinking Mapper—and then invoke diffusion priors guided by the CLIP embedding, a coarse EEG-derived image, and auto-captions to stabilize generation, achieving state-of-the-art zero-/few-shot performance and indicating that informative semantic structure emerges around 250 ms post-stimulus (Li et al. 2024; Craik et al. 2019; Bai et al. 2023; Song et al. 2023). MEG follows a similar strategy: neural activity is aligned to pretrained visual embeddings (CLIP, DINOv2) through regression or contrastive learning, and a diffusion model is subsequently conditioned to reconstruct images; time-resolved analyses reveal peaks that track high-level semantics and outperform linear decoders by large margins in retrieval settings (Benchetrit et al. 2023).

Extending from static images to video requires addressing the temporal mismatch between second-scale hemodynamics and high frame rates. Recent fMRI systems therefore separate *semantics* from *perception*: windowed BOLD signals are mapped to CLIP-like keyframe embeddings while a complementary pathway estimates low-level appearance and motion flow; both streams condition a pretrained text-to-video diffusion model to yield temporally coherent clips with improved SSIM and cross-frame consistency (Gong et al. 2024; Chen et al. 2023b; Zhu et al. 2025; Sun et al. 2024).

Overall, fMRI remains strongest for spatial detail, whereas EEG and MEG excel at temporal dynamics; FM-based alignment (e.g., CLIP) and generative priors (diffusion) compensate for each modality’s weaknesses and can be adapted with minimal architectural changes across recording types (Liu et al. 2025a; Craik et al. 2019; Li et al. 2024). At the same time, the degree to which this “neural-to-semantic alignment + conditional diffusion generation” recipe generalizes beyond controlled benchmarks and subject-specific settings is still inconsistently demonstrated.

#### Failure modes & boundary conditions

FM-conditioned reconstruction often depends on accurate subject-specific mapping into a shared semantic latent, which may degrade under session drift, noise, or mismatched preprocessing. Diffusion priors can yield visually plausible images even when neural evidence is weak, creating a perceptual-quality vs neural-fidelity gap. Cross-dataset transfer is often fragile because voxel/channel selection, stimulus distributions, and evaluation pipelines differ.

#### Contradictions across studies

Reported gains are not always comparable because papers optimize different targets and use different conditioning signals and sampling settings. Methods emphasizing semantic alignment may trade off fine-grained structural fidelity, and “State-of-the-art” claims can be confounded by within-subject vs cross-subject protocols. Standardized metrics and splits are therefore necessary to interpret improvements consistently.

#### Reliability & reproducibility signals

Evidence is concentrated on a few canonical benchmarks (e.g., NSD/GOD summarized in Table 4), which aids comparability but risks overfitting to shared routines. Reliability is stronger when multi-subject and cross-session results are reported with clear pre-processing and split details, yet these signals are uneven. We recommend routine reporting of subject/session counts, split strategies, and both fidelity- and semantics-oriented metrics.

### 5.3 Decoding Language and Speech

Recent progress in non-invasive language and speech decoding has come from treating foundation models (FMs)—large language models (LLMs) and self-supervised sequence/speech encoders—as priors, targets, and generators within end-to-end pipelines (overview in Table 5; schematic in Figure 4) (Sato and Kobayashi 2025; Chen et al. 2024b; Qiu et al. 2025b). The central idea is to align neural activity to pretrained semantic or acoustic manifolds and then exploit powerful generative or retrieval mechanisms to recover continuous content.

Despite fMRI’s slow temporal dynamics, LLM-centric pipelines enable continuous semantic decoding by using the language model as a prior/decoder to enforce coherence (Sato and Kobayashi 2025). A common recipe builds an encoding model that predicts fMRI responses from linguistic features (Sun et al. 2020) and then scores candidate continuations sampled from an LLM (e.g., GPT), selecting the sequence that best matches the predicted brain activity—trading data for fluency and substantially mitigating temporal limits (Tang et al. 2023). Recent research has further scrutinized this alignment, revealing that while neural language models are powerful, psychologically plausible models can sometimes offer superior alignment with brain representations in specific linguistic contexts (Zhang et al. 2024). Performance scales with model size but typically requires sizable per-subject data; nonetheless, systems can recover the gist of perceived/imagined narratives or silent videos, indicating access to higher-level semantics beyond low-level cues (Tang et al. 2023; Antonello et al. 2023). For example: *MindLLM* targets subject-agnostic decoding via a neuroscience-informed attention encoder plus “Brain Instruction Tuning” with an off-the-shelf LLM (Qiu et al. 2025a); *UMBRAE* aligns fMRI into a VLM latent space to flexibly decode text and/or images (Xia et al. 2024); and *GPT-2 prompting* directly treats fMRI as prompts aligned to text prompts through contrastive learning (Chen et al. 2024b). Early language decoding work primarily focused on lexical-level decoding. Subsequently, research began to shift towards more complex sentence-level semantic representations (Sun et al. 2019). Overall, FMs supply powerful linguistic constraints and generation, advancing versatility and cross-subject generalization while compensating for fMRI’s temporal limitations (Sato and Kobayashi 2025; Tang et al. 2023; Qiu et al. 2025a).

EEG’s millisecond precision shifts the focus to acoustic dynamics. Pipelines map EEG into pretrained speech representation spaces (e.g., Wav2Vec 2.0) using regression or contrastive objectives, after which CTC/sequence decoders or retrieval heads infer phonemes and words; this transfer learning improves robustness with limited labels and supports cross-subject generalization (Craik et al. 2019; Qiu et al. 2025b). The approach leverages FM encoders as invariant acoustic priors while allowing lightweight adaptation to individual subjects and sessions.

MEG occupies a complementary sweet spot: its temporal fidelity enables fine-grained analyses while offering better source separability than EEG. Neuroscience-inspired self-supervised learning on unlabeled MEG (masking, band prediction) yields representations that scale with data and generalize across tasks and subjects, setting strong baselines for phoneme/word decoding under data scarcity (Jayalath et al. 2024). Aligning MEG features to pretrained speech models further boosts recognition of perceived speech and supports time-resolved probing of linguistic feature dynamics (Jayalath et al. 2024; Qiu et al. 2025b).

Looking toward practical communication BCIs, non-invasive routes based on fMRI (Tang et al. 2023), potentially fNIRS, and EEG spellers augmented with FM priors (Carìa 2025) show growing promise but face constraints in equipment bulk, calibration demands, latency, and robustness. Across modalities, the FM recipe—neural-to-semantic/acoustic alignment plus generative or retrieval decoding—often enables richer outputs from weak signals, yet robust cross-session and cross-subject performance remains unevenly validated.

#### Failure modes & boundary conditions

Language/speech pipelines are sensitive to intermediate targets (tokens vs embeddings vs acoustics) and to the quality of brain-to-linguistic mapping. Injecting brain-derived conditions into frozen LLMs can be brittle when the condition signal is noisy, especially for open-vocabulary generation, while end-to-end routes often require more paired data and careful regularization. For imagined-speech EEG, low SNR and inter-session variability frequently dominate performance.

#### Contradictions across studies

Results diverge because evaluation emphasizes different endpoints (WER/CER, semantic similarity or perceived intelligibility) and different constraints (closed vocabulary/spellers and open-ended generation). Protocols also vary in whether candidate reranking (AG-EML) is used, which changes task difficulty. Without standardized splits and explicit decoding constraints, cross-paper ranking remains partially confounded.

#### Reliability & reproducibility signals

Reliability is higher when multi-subject and cross-session validation is reported together with fully specified preprocessing and decoding hyperparameters. However, datasets and reporting practices remain heterogeneous (as reflected in Table 5), and code availability is uneven. We recommend documenting subject/session counts, split strategies, and paired reporting of literal fidelity and semantic consistency metrics.

### 5.4 Decoding Audio, Music, and Auditory Attention (AAD)

Decoding auditory experiences—ranging from attended speech to complex music—has benefited from foundation-model (FM) ideas, though progress lags behind vision and language (Park et al. 2023). A central thread is auditory attention decoding (AAD), which aims to infer the attended source (e.g., speaker) from non-invasive brain signals to enable neuro-steered hearing aids (Geirnaert et al. 2021). Early linear pipelines (stimulus/temporal response functions, CCA) (De Taillez et al. 2020) provided a principled starting point (Wong et al. 2018; Fiedler et al. 2017), but they struggle with nonlinearity and short decision windows that are critical for real-time control (Puffay et al. 2023; Cai et al. 2020; Vandecappelle et al. 2021). Recent systems replace these with non-linear encoders that borrow FM-inspired components—self-attention, convolutional hierarchies (Fan et al. 2024; Su et al. 2021; Xu et al. 2022; Accou et al. 2023), and graph neural networks—to model spatio-temporal structure and intersensor topology, yielding markedly higher accuracy at short latencies (Cai et al. 2023; Fan et al. 2025b,a; Li et al. 2025b). Within this family, some approaches perform identity-based AAD by correlating EEG with reference speech streams using CNN/attention encoders or encoder–decoders (Su et al. 2021; Xu et al. 2022; Accou et al. 2023); others pursue reference-free spatial AAD by estimating the attended location directly from EEG via spatio-temporal attention, EEG-graph networks, dynamic graph self-distillation networks, or recent hybrid/mamba-style architectures (Accou et al. 2023; Fan et al. 2024; Zhou et al. 2025; Ni et al. 2024; Fan et al. 2025b; Yan et al. 2024; Fan et al. 2025a). Collectively, these non-linear models narrow the gap to practical neurosteered hearing aids by improving robustness in short windows (Geirnaert et al. 2021; Zhang et al. 2023).

Beyond attention, reconstruction of auditory content is emerging. For fMRI, hierarchical feature decoding leverages knowledge of the auditory pathway: models first decode deep-network–inspired auditory features from BOLD and only then synthesize waveforms using an audio generative model, thereby avoiding a direct slow-fMRI-to-fast-audio mapping (Park et al. 2023). For EEG, millisecond temporal precision suits rhythm and envelope tracking; recent work employs latent diffusion models for naturalistic music reconstruction, pushing beyond classical linear envelope decoding though typical quality remains below speech reconstruction (Postolache et al. 2025; Simon et al. 2022). Notably, DMF2Mel (Fan et al. 2025c) has achieved precise reconstruction of minute-level continuous imagined speech, effectively balancing temporal dependency modeling and information retention efficiency in long-sequence decoding. These results underline both the promise of generative priors and the challenge of capturing rich musical structure from sparse, noisy measurements.

Comparative analyses highlight systematic differences between music and speech decoding. Linear EEG models reveal shared basic acoustic tracking yet lower reconstruction quality for music envelopes, consistent with distinct cortical processing pathways and with the added structural complexity of music (Simon et al. 2022). This complexity is being used as a probe for auditory scene analysis, while fMRI studies map genre-level representations across cortex (Nakai et al. 2021). Overall, FM-inspired encoders and generative decoders provide a coherent route for AAD and auditory reconstruction: learn invariant, topology-aware neural representations; align them with pretrained acoustic manifolds; and leverage strong priors for synthesis—steps that collectively improve short-window accuracy for attention control and expand the scope of non-invasive reconstruction of sound and music. However, evidence under realistic acoustic mixtures, strict latency constraints, and cross-session robustness is still comparatively limited.

#### Failure modes & boundary conditions

AAD is constrained by decision-window length, acoustic scene complexity, and session variability; gains under controlled mixtures may not persist under realistic noise and artifacts. For auditory reconstruction, strong generative priors can mask weak neural evidence, and capturing long-range musical structure remains difficult from noisy measurements. In hearing-aid-like deployments, latency and calibration stability are often the bottlenecks.

#### Contradictions across studies

Comparisons differ due to stimulus type (speech and music), availability of clean references, window definitions, and metric choices (accuracy and correlation-based scores). Deep/FM-inspired models can show large gains in some controlled settings, while classical TRF/CCA baselines remain competitive when data are limited or interpretability and stable calibration matter. This motivates matched baselines and standardized protocols.

#### Reliability & reproducibility signals

Reliability improves when studies report cross-session validation, realistic mixtures, and strong matched baselines under consistent preprocessing. Reporting of attention labels, windowing, and latency/compute constraints is still inconsistent, limiting reproducibility. We recommend standardizing window lengths, reporting latency, and including both classical and FM-based baselines.

### 5.5 Multimodal Decoding

A growing line of work moves beyond single-modality pipelines to fuse complementary signals—across neural modalities (e.g., fMRI+MEG/EEG, EEG+fNIRS) and with auxiliary behavioral streams (e.g., eye tracking)—while aligning all channels to shared FM latents (CLIP/VLM or speech encoders) for generation or retrieval. Methodologically, these systems pursue two converging threads: (i) neural–neural fusion to combine fMRI’s spatial precision with MEG/EEG’s millisecond dynamics or fNIRS’s portability, and (ii) neural–semantic alignment that maps each input stream into a unified FM-conditioned space where decoding and cross-modal transfer are performed with minimal architectural changes.

On the neural–neural side, unified frameworks such as UMBRAE learn a shared latent in which different brain measurements (and even different stimulus types) can be decoded to image or text outputs; contrastive and generative objectives encourage instance-level alignment and enable flexible “modality in, task out” decoding (Xia et al. 2024). Complementary work explicitly aligns EEG, MEG, and fMRI representations in a common space to support decoding, encoding, and modality conversion, showing that an FM-style alignment layer can bridge disparate temporal/spatial scales and improve cross-subject generalization (Ferrante et al. 2024c). At the dataset level, recent large, paired M/EEG–fMRI resources and fusion analyses demonstrate consistent gains in modeling naturalistic visual processing when joint information is exploited, reinforcing the value of multi-neural supervision for both training and evaluation (McNabb et al. 2025; Wang et al. 2022b). In portable settings, hybrid EEG+fNIRS architectures with cross-modal attention or graph fusion markedly outperform unimodal baselines on cognitive/affective tasks, suggesting a practical path toward out-of-lab BCIs that still benefit from FM-style invariances (Li et al. 2025c).

For neural–semantic alignment, multimodal decoders pair fused neural embeddings with strong generative priors. Visual decoders condition diffusion backbones on a joint latent that aggregates, for example, fMRI-derived spatial codes with MEG/EEG temporal embeddings, improving semantic retrieval and frame consistency for naturalistic stimuli; joint video–text embeddings further enhance encoding/decoding of movie stimuli versus uni-modal feature baselines (Fu et al. 2025b). Auxiliary behavioral streams can be folded in as well: MR-based eye tracking (DeepMReye) provides gaze priors directly from fMRI volumes, offering an inexpensive behavioral channel that stabilizes reconstructions and disambiguates scene context when combined with neural features (Frey et al. 2021). Together, these designs operationalize a general recipe: map each stream (fMRI/MEG/EEG/fNIRS/behavior) into a shared FM-conditioned latent, perform attention or contrastive fusion with modality-dropout for robustness, and decode via conditional diffusion or VLM heads.

Looking ahead, multimodal decoding raises new opportunities and open problems: scalable co-training across sites and sensors; modality-adaptive schedulers that weight channels by SNR and task phase; and foundation models for neural data that natively support modality conversion (e.g., fMRI to MEG) and few-shot personalization. Early evidence from unified decoders and aligned representations indicates that fusing complementary signals consistently boosts semantic fidelity and temporal precision, delivering a coherent blueprint for FM-driven multimodal brain decoding in both lab-grade and portable settings.

### 5.6 Interpreting Metrics across Research

Tables 4 and 5 summarize the metrics reported in recent visual and language/speech decoding research. However, a key reason why quantitative synthesis remains difficult is that these metrics correspond to different evaluation objectives and are often calculated under incompatible protocols; therefore, the same metric names do not necessarily allow for comparison and support of the same conclusions.

We categorize the metrics into three types. The first is **fidelity**, which examines whether the decoded output is supported by neural evidence or matches explicit ground truth (e.g., SSIM / PSNR, PixCorr / PCC, N-way Accuracy for images; WER / CER/HER for speech under specific decoding constraints). The second is **perceptual quality** (e.g., FID / IS, Human Accuracy), which reflects realism or human subjective judgment or preference. However, it may be influenced by generative priors and therefore cannot serve as independent evidence for decoding evaluation. The third is **semantic consistency** (e.g., PSM / ROUGE / BLEU / METEOR / CIDEr / SPICE or BERTScore), which assesses meaning preservation. These three types of metrics can complement each other but are not interchangeable; even if fidelity decreases, perceptual metrics may improve. Strict pixel or n-gram overlap scores may penalize semantically correct outputs in cases of paraphrasing or appearance changes.

Incompatibility between different research findings also stems from data acquisition methods, candidate set construction for identification, preprocessing and channel count, feature extraction and normalization, and cueing and decoding settings. Therefore, interpretation should only be based on the reported protocol. We recommend that research reports validate the protocol, along with at least one fidelity-oriented metric and one semantic or utility-oriented metric. Key protocol details required for reproducible scoring should also be disclosed (e.g., grouping design, candidate set, and decoding/generation settings). Cross-paper comparisons should be limited to studies employing the same protocol and evaluating the same target attribute. Quantitative ranking between papers is only meaningful within a subset that employs the same protocol and consistently evaluates the target attribute.

## 6 Cross-Domain Trends and Breakthroughs

Across vision, language, and audio/speech decoding, pre-trained foundation models (FMs)—diffusion generators, CLIP-style dual encoders, large language models (LLMs), and self-supervised Transformers—have become pervasive as feature extractors, priors, and generators (Gong et al. 2024; Sato and Kobayashi 2025; Park et al. 2023; Jayalath et al. 2024). A unifying blueprint is to first align brain activity with an FM latent space and then leverage the FM’s generative or retrieval capacity for reconstruction or recognition. Alignment is achieved by (i) contrastive learning that places neural and stimulus embeddings in a shared manifold (Li et al. 2024; Chen et al. 2024b), (ii) direct regression that predicts FM embeddings or conditioning variables from neural signals, or (iii) “encoding-model inversion,” where a brain-prediction model guides an LLM-based search to select sequences whose predicted neural responses best match observed data (Tang et al. 2023).

Within this framework, FMs systematically compensate for modality-specific weaknesses. For fMRI, LLM priors impose linguistic structure over slow hemodynamics, enabling coherent semantic decoding, while keyframe-plus-flow strategies reduce temporal aliasing in video reconstruction (Tang et al. 2023; Gong et al. 2024). For EEG/MEG, strong visual priors from CLIP and diffusion models elevate semantic image reconstruction despite limited spatial precision and low SNR, and self-supervised learning extracts robust time-resolved features that generalize across tasks and subjects (Li et al. 2024; Jayalath et al. 2024). These advances coincide with a clear shift toward subject-agnostic and zero/few-shot regimes—e.g., MindLLM, SSL-trained MEG encoders, and zero-shot EEG visual decoding—aimed at reducing per-subject calibration and improving real-world viability (Qiu et al. 2025a; Jayalath et al. 2024; Li et al. 2024; Tang et al. 2023).

A second trajectory is deeper multimodality: systems increasingly align brain activity to vision-language models (VLMs), enabling flexible decoding into text and/or images within a single framework (e.g., UMBRAE) (Qiu et al. 2025a; Xia et al. 2024). In parallel, the community is probing temporal dynamics more explicitly in EEG/MEG, using FM-based analyses to characterize real-time processing, feature persistence, and semantic stabilization windows (Benchetrit et al. 2023; Li et al. 2024; Jayalath et al. 2024). Finally, generative pipelines themselves are becoming more sophisticated—multi-stage designs, multi-feature conditioning, and specialized architectures now dominate SOTA reconstructions—reflecting a maturation from simple stimulus-to-signal mappings to modular, prior-driven generation (Gong et al. 2024; Li et al. 2024; Chen et al. 2023b).

## 7 Persisting Challenges and Future Horizons

Despite remarkable progress in leveraging foundation models (FMs) for brain-computer interfaces (BCIs), several major obstacles persist that limit real-world deployment, particularly in decoding speech and visual experiences from brain signals. This section integrates challenges and opportunities across modalities.

### 7.1 Current Hurdles for FMs in BCI

Despite rapid progress, foundation-model–based decoding faces several structural obstacles. First are computational and data constraints. State-of-the-art FMs demand substantial training and inference resources that are difficult to accommodate in wearable or clinical settings; latency and power budgets limit real-time use on embedded hardware. At the same time, effective pretraining and reliable evaluation require large, diverse, and well-annotated datasets, which are costly and slow to acquire in neuroscience. Speech tasks often lack sufficient labeled EEG/MEG, and visual decoding likewise depends on expensive high-resolution fMRI or specialized corpora (e.g., NSD, THINGS), constraining both model scaling and generalization (Chen et al. 2023a; Jiang et al. 2024; Shirakawa et al. 2024). These constraints also shape the strength of evidence available in current FM-based decoding studies. We therefore distinguish four validation regimes with increasing translational relevance: within-session, single-subject evaluation; cross-session evaluation on the same subject (to probe drift); cross-subject evaluation (to assess population-level generalization); and naturalistic or online settings with latency and robustness constraints. Much of the literature remains concentrated in the first two regimes, whereas reproducible gains in cross-subject and deployment-oriented settings are less consistently demonstrated. This distinction is useful for interpreting “generalization” claims throughout this section.

A second hurdle arises from signal quality and modality trade-offs. EEG/MEG offer millisecond timing but suffer from low spatial specificity, non-stationarity, and vulnerability to artifacts; reliable access to deep sources is limited relative to fMRI. Conversely, fMRI localizes fine-grained spatial structure but blurs rapid dynamics due to slow hemodynamics. Managing physiological and environmental noise—especially in EEG—remains non-trivial, and mapping complex stimuli such as natural language onto measured neural responses is rarely one-to-one, complicating model design and evaluation (da Silva 2013; Logothetis 2008).

Inter-subject variability further impedes robustness. Anatomical differences, functional topographies (e.g., retinotopy), state fluctuations, and session effects degrade cross-subject transfer; models trained on one individual often fail to generalize to others (i.e., Level-3 cross-subject validation) without costly calibration. Achieving zero-shot or strongly subject-agnostic performance via functional alignment or adaptation remains an open challenge across language and vision decoding alike (Luo et al. 2024; Chen et al. 2023a).

Model complexity and decoding ambiguity compound these difficulties. Open-vocabulary generation (free-form text or speech) is markedly harder than closed-set recognition, particularly for EEG. Higher-level semantic decoding is less mature than phonetic decoding, and imagined/silent speech (or internally generated imagery such as dreams) presents weaker, noisier signals without clear behavioral anchors. Architectures must also respect modality structure (e.g., irregular sensor graphs for EEG, 3D sampling for fMRI), while delivering low-latency, reliable outputs suitable for interactive BCIs which becomes most stringent under Level-4 online or naturalistic constraints. In parallel, debates persist about the cognitive plausibility of very large FMs as models of human processing and how best to constrain them for neuroscientific validity.

In addition, interpretability and evaluation remain limited. Many pipelines behave as black boxes, with unclear spatiotemporal mappings from brain activity to outputs; linear probes and attention maps provide partial insight but fall short of comprehensive explanations. Standard metrics (pixel-wise scores for images, BLEU/WER for language) can poorly reflect perceptual or semantic fidelity, underscoring the need for human-centered and task-relevant evaluations (Gong et al. 2024).

Consequently, the black-box opaqueness of these models exposes a vulnerability to targeted adversarial attacks. Many recent systems rely on an alignment step which maps brain signals into a high-dimensional latent space, used for retrieval or conditional generation. This acts as an attack surface where adversarial perturbations can be crafted to induce small yet semantically meaningful shifts in the representation, altering retrieval outcomes or the conditioning signals passed to a generator (Zhang et al. 2020; Lubars and Chin 2022; Li et al. 2021). As generative models are able to produce fluent text and images given weak or ambiguous input, this induced misalignment often manifests as a high-confidence but semantically incorrect - hallucinated - output rather than an obvious corruption (Ji et al. 2023). As discussed earlier, risks of both test-time evasions and training-time backdoors are established threats in the literature (Meng et al. 2022). Therefore, robustness claims for FM-based decoders must explicitly state the assumed threat model and report rigorous adversarial stress-testing. Future work to quantify attack success rates under evasion and adversarial perturbations, alongside basic backdoor checks, would be recommended to enhance standard denoising pipelines to account for intentional, alongside accidental, noise.

At the same time, reproducibility and reporting are considerable bottlenecks in non-invasive brain decoding. In non-invasive brain decoding, claims are only as reproducible as the specificity with which the full experimental and computational pipeline is documented. Authors should therefore make the data provenance and cohort structure explicit (dataset release or acquisition protocol, access/consent constraints, subject and session counts, inclusion/exclusion rules, and a precise task specification with timing and label definitions), and provide a technically complete description of acquisition and preprocessing (instrumentation, sampling rate/TR, channel/voxel specification and referencing, followed by the exact sequence of denoising, artifact handling, normalization/alignment, and any modality-specific steps such as fMRI motion correction, together with ROI/voxel/channel selection and time-windowing). Equally important is a split protocol described at a level that rules out leakage (within-subject, cross-session, or cross-subject, with fold construction stated), since this choice often dominates reported performance; these split choices correspond directly to the validation regimes above and should be treated as part of the primary evidence claim. Method reporting should not stop at model names: the FM backbone version, alignment target space, optimization setup, and random seeds must be specified; when a generative component is used, prompts and decoding settings should be treated as part of the method and reported accordingly. Finally, evaluation should define metrics in relation to the quantity of interest (fidelity versus perceptual quality versus semantic consistency) and, where feasible, provide uncertainty estimates; releasing code, pre-processing scripts, configurations, and checkpoints remains the most direct mechanism for enabling independent verification.

Finally, ethical considerations are central. Brain data are intrinsically sensitive, demanding stringent privacy, security, and governance to address ’mind-reading’ concerns and clarify ownership. FM training can amplify demographic and contextual biases, raising equity issues in performance and access. Recent neuro-rights proposals argue that neural data processing implicates rights such as mental privacy and cognitive liberty (Ienca and Andorno 2017; Yuste et al. 2017), motivating protections beyond conventional biomedical data governance (European Parliament 2024; UNESCO 2023). This challenges the adequacy of a static consent system where a single, one-off signature is obtained. As alignment techniques, prompting interfaces and downstream reuse evolve, decoders may enable secondary inferences beyond the user’s original intent (F. et al. 2019; Future of Privacy Forum and IBM 2021) such as biometric ’brainprints’ (Yang et al. 2022), mood/intent inference, or other latent attributes. A practical path forward is adoption of a tiered consent system which explicitly separates permissible classes of outputs/inferences (e.g., task labels vs. semantic reconstructions vs. affect/clinical inference), alongside dynamic consent mechanisms which support ongoing re-consent, granular preference updates, withdrawal and audit for when the use cases change over time (Kaye et al. 2015; Budin-Ljøsne et al. 2017; Wiertz and Boldt 2022).

### 7.2 Applicability Boundaries for FM-Based BCI

Foundation Model brain decoding is usually built around two pieces: a learned mapping from brain activity into an intermediate representational space, and a strong generative prior that turns that representation into a perceptible output. This design can noticeably improve output richness in controlled experiments, but its reliability is uneven once you move across subjects, across sessions, or into settings with tight deployment constraints. Performance often drops when the alignment itself drifts (e.g., day-to-day or subject-to-subject variability), when the generator “fills in” missing evidence with plausible content, or when results hinge on prompting choices and decoding hyperparameters. Taken together, these limitations argue for stating the operating boundaries explicitly, rather than treating FM-based pipelines as a universal upgrade.

Classical pipelines remain the better default in several high-value regimes. For low-latency, closed-set control tasks (P300/SSVEP/MI), lightweight classifiers paired with well-understood signal-processing steps are typically more stable and easier to deploy than generative decoding. When data are scarce and drift is unavoidable, sample-efficient and interpretable baselines, such as TRF/mTRF and CCA in auditory attention decoding—are often easier to regularize and to validate under cross-session evaluation. In MI classification, covariance-based Riemannian methods have a long track record of robust performance with modest data requirements (See Table 6 for details). Across these cases, the priority is reliability, calibration stability, and latency; adding large generative components can increase complexity without delivering consistent gains.

The same ingredients that make FM-based pipelines attractive can also make failure modes sharper. If the brain-to-semantic mapping is even slightly misaligned, the downstream generator may amplify that error into a confident-looking output. Moreover, a strong prior can mask weak neural evidence, producing results that score well on perceptual criteria yet deviate from the underlying signal—especially when evaluation emphasizes plausibility over neural fidelity. Finally, both LLM- and diffusion-based modules are sensitive to prompting and decoding settings, which can introduce avoidable variance unless standardized. For these reasons, FM-based improvements should be demonstrated under matched latency budgets, explicit cross-session and cross-subject splits, and a metric suite that jointly captures fidelity and semantic consistency.

### 7.3 Opportunities and Future Research Directions

Meeting today’s challenges points to a clear research agenda. A first priority is the development of Brain Foundation Models (BFMs): large, modality-aware encoders pretrained on diverse fMRI/EEG/MEG corpora to learn universal neural representations. Progress will hinge on self-supervised learning tailored to neural signals (masking, predictive objectives, cross-modal contrastive alignment), on hybrid architectures that bridge modalities (e.g., EEG to Text or EEG to Audio contrastive models), and on theoretically informed inductive biases such as predictive coding. Two complementary threads should run in parallel: systematically probing how scale, objective choice, and adaptation strategy affect encoding/decoding performance and cognitive plausibility; and leveraging powerful existing FMs (speech recognizers, LLMs, VLMs) as computational probes of brain representations.

A second frontier is decoding internally generated content. In speech, this includes silent speech and inner monologue; in vision, mental imagery and dreams. Because such phenomena lack overt behavioral anchors and unfold with weak, distributed signals, they will likely require new representational targets (semantic, affective, or episodic manifolds) and training curricula grounded in cognitive neuroscience—e.g., predictive coding or generative models that explicitly separate latent intent from sensory realization.

Third, advancing robustness and real-time BCIs remains essential. Methods must better tolerate noise, artifacts, and non-stationarity, while generalizing across subjects, tasks, datasets, and even modalities via functional alignment and adaptation. For EEG/MEG, rigorous comparisons between sensor-space and source-reconstructed decoding are needed; for open-vocabulary language and rich visual content, newer generative backbones may improve fidelity, controllability, and latency. The overarching goal is reliable, low-latency decoding suitable for interactive use in realistic settings.

Clinical translation is the fourth axis. Communication BCIs for individuals with ALS, stroke, or paralysis are a primary driver; success depends on robust models, streamlined calibration, and user-centered design (comfort, intuitive interfaces), alongside regulatory-grade reliability. Beyond assistive communication, decoding pipelines can serve basic neuroscience—clarifying language processing, semantic representation, and cognitive dynamics—and may yield biomarkers for neurological and psychiatric disorders. Emerging modalities such as HD-DOT add promising trade-offs for naturalistic language decoding and should be systematically evaluated.

Achieving these aims will also require efficient model design and innovative data strategies. Compression, distillation, parameter-efficient fine-tuning, and compact architectures can bring state-of-the-art decoding to edge and wearable hardware. On the data side, priorities include building large public datasets (e.g., via OpenNeuro), advancing augmentation for neurodata, expanding SSL on unlabeled recordings (Jayalath et al. 2024), and adopting privacy-preserving collection and training (federated/secure aggregation) to diversify cohorts without compromising confidentiality.

Generalization and personalization form a sixth pillar. Domain adaptation, meta-learning, and few-shot calibration can minimize per-subject effort; online adaptation can track drift; and cross-modal transfer can bootstrap under-resourced settings. Training on naturalistic data—continuous narratives, real-world scenes—should become standard to narrow the lab-to-life gap.

Interpretability and governance complete the road-map. XAI for BCI—from attention maps and feature attributions to causal perturbations and encoding-model probes—can validate neuroscientific plausibility, surface spurious shortcuts, and build user trust. Bridging the lab-to-life gap further requires durable hardware, stable signal quality over months, and algorithms robust to long-term drift, all under ethical frameworks that foreground privacy, consent, equity, transparency, and accountability. Finally, treating FMs as tools for neuroscience—to instantiate and test hypotheses about neural information processing—offers a deep synergy: decoding improves as our models of the brain improve, and vice versa (Huth et al. 2016). In the long term, reliable decoding of inner speech and abstract thought remains a moonshot goal, one that will likely emerge from the convergence of BFMs, theory-guided objectives, efficient adaptation, and ethically grounded, human-centered design (Tang et al. 2023).

This roadmap is based on the premise that evaluation results in this field are comparable across different studies and meaningful for real-world deployment. Currently, many studies obtain results under different experimental protocols, such as different subjects, preprocessing procedures, and decoding constraints. Furthermore, the metrics used also vary, thus failing to reflect the same underlying conclusions (Jayaram and Barachant 2018). This is crucial for practical applications in brain decoding; only through explicit cross-session and cross-subject validation can robustness to drift, low-latency performance, and calibration burden be reliably assessed (Lotte et al. 2007). Ideally, stress tests simulating natural conditions should also be conducted. Therefore, we recommend and advocate for the use of a minimal benchmarking protocol for future FM-based BCI research: 1) report at least one cross-session evaluation and, where feasible, one cross-subject evaluation; 2) compare against a highly matched baseline under the same preprocessing conditions (Glaser et al. 2019); 3) document the cueing and decoding/generation settings as part of the methodology; and 4) use a set of metrics that distinguishes between fidelity, semantic consistency, and task usefulness, while reporting uncertainty. In addition to making different studies comparable, this standardized benchmarking scheme also lays the foundation for subsequent discussions on deployment and dual-use, making empirical capabilities more clearly defined. For clarity, Table 7 provides a prioritized roadmap connecting major obstacles with research objective, actionable methodology and reference benchmarks. This roadmap emphasizes that comparability (P1) is a prerequisite for assessing robustness (P2) and deployment feasibility (P3), while also regarding credible decoding and governance (P4) as equally important objectives, rather than as ex-post considerations. These directions collectively put the evidence hierarchy discussed above into practice, clarifying the assessment mechanisms and reporting expectations.

Against this backdrop, while neural decoding holds clear clinical promise such as in communication interfaces (Anumanchipalli et al. 2019; Willett et al. 2021a) and rehabilitation in severe motor impairment (Chaudhary et al. 2016), the same representational power can be repurposed for non-therapeutic and potentially coercive uses. This creates a dual-use dilemma where advances that improve decoding fidelity, generalisation and robustness also increase the plausibility of surveillance applications (Zuboff 2019). This includes attempts to infer sensitive mental content from non-consenting subjects, as well as workplace and consumer monitoring (Future of Privacy Forum and IBM 2021) and ’neuromarketing’-style behavioural optimisation. These capabilities in strategic contexts raise questions about military and intelligence adoption pathways even when current systems remain technically constrained (Claverie and du Cluzel 2022; Defense Advanced Research Projects Agency (DARPA) 2018). From a research perspective, this motivates treating misuse-resistance as a priority objective alongside accuracy. Concrete directions include explicit threat modelling and red-teaming for coercive or non-consensual settings, governance and deployment constraints such as purpose limitation, access control and secure logging (European Parliament 2024; UNESCO 2023). Finally, the use of technical safeguards such as encryption and differential privacy which can reduce unintended inference and unauthorized reuse across diverse tasks or contexts (Bonaci et al. 2014; Future of Privacy Forum and IBM 2021).

## 8 Conclusion

Foundation models have substantially accelerated progress in non-invasive brain decoding by providing transferable representations and strong generative priors that improve semantic alignment and reconstruction quality across modalities. By leveraging large-scale pre-training, transfer learning, and powerful generative capabilities, FMs demonstrate an unprecedented ability to interpret complex patterns in noisy signals from EEG, MEG, fMRI, and fNIRS. This survey highlighted breakthroughs: high-fidelity visual reconstruction (images/video) across modalities using CLIP and diffusion models; continuous semantic language decoding from fMRI via LLMs overcoming temporal limits, with progress towards subject-agnostic models; enhanced AAD performance using FM-inspired architectures; and emerging success in reconstructing complex sounds and music using generative approaches. Collectively, these advances broaden what is achievable in controlled settings, while robust cross-subject generalization and deployment-level reliability remain open challenges.

However, the journey to widespread, reliable, and ethical deployment remains challenging. Computational/data requirements, interpretability needs, robust generalization (across subjects, tasks, environments), and profound ethical considerations persist. Continued innovation is needed not just in scaling models, but in developing efficient, adaptable, explainable, and personalized approaches tailored to BCI constraints.

The future lies at the intersection of AI, neuroscience, engineering, and ethics. Realizing FM’s full potential requires parallel progress in sensing hardware, diverse datasets, realistic validation benchmarks, and proactive ethical dialogue. The ultimate goal is to harness FMs not only for powerful decoding tools but also to deepen our understanding of the brain, ensuring responsible development for human benefit. Continued interdisciplinary collaboration is paramount in transforming the promise of FM-driven non-invasive brain decoding into tangible improvements in health, communication, and human-world interaction.

In summary, FM-enabled decoding represents a compelling path toward assistive communication (Anumanchipalli et al. 2019; Willett et al. 2021a) and rehabilitation (Chaudhary et al. 2016), but its societal impact will depend on how capability is governed and constrained. As models become more general and decoding outputs more semantically expressive, the boundary between clinical assistive use and surveillance-oriented inference can narrow. This risk is particularly acute when neural data is linked to identity and used beyond the original task’s context (Future of Privacy Forum and IBM 2021), highlighting the need for robust institutional oversight (European Parliament 2024; UNESCO 2023). A mature research agenda should therefore explicitly acknowledge the dual-use potential and incorporate safety-by-design principles accordingly. Technical progress must be aligned with protections for mental privacy and cognitive liberty (Yuste et al. 2017; European Parliament 2024; UNESCO 2023), ensuring that sufficient technical safeguards are embedded with auditable accountability mechanisms from the development phase (Bonaci et al. 2014).

## Acknowledgements

We acknowledge the support from the UMRI IDR Placement 2025 Pioneering project ”Foundational AI Models for Brain Computer Interfaces and Neuro-Robotic Control”, and The European High Performance Computing Joint Undertaking (EuroHPC JU) with project EHPC-DEV-2025D06-002 and project EHPC-BEN-2025B05-008.

## Declarations

### Funding

This work is supported by the STI 2030—Major Projects (No. 2021ZD0201500), the National Natural Science Foundation of China (NSFC) (No.62571002, 62406058), Excellent Youth Foundation of Anhui Scientific Committee (No. 2408085Y034); Sichuan Science and Technology Program (2025ZNS-FSC1480); China Postdoctoral Science Foundation (2024M750360); Fundamental Research Funds for the Central Universities (ZYGX2024XJ051). Natural Science Foundation of China (62472206), National Key R&D Program of China (2025YFC3410000), Shenzhen Science and Technology Innovation Committee (RCYX202312 11090405003, KJZD20230923115221044).

### Conflict of interest

The authors have no relevant financial or non-financial interests to disclose.

### Ethics approval and consent to participate

Not applicable. This article is a review/survey study and does not involve new studies with human participants or animals performed by any of the authors.

### Consent for publication

Not applicable.

### Data availability

No new data were generated or analysed in this study. This article is a survey of previously published literature, and dataset information summarized in Table 2 is derived from the cited sources.

### Materials availability

Not applicable.

### Code availability

No code was generated or used in this study.

### Author contribution

Yifan Wang: Data curation, Investigation, Methodology, Visualization, Writing – original draft, Writing – review and editing. Shaonan Wang: Conceptualization, Supervision, Writing – original draft, Writing – review and editing. Wenhao Cai: Visualization. George Ford: Writing – review and editing. Yang Cui: Writing – review and editing. Yunhao Zhang: Methodology, Writing – original draft. Changde Du: Writing – original draft. Cunhang Fan: Funding acquisition, Writing – original draft. Dongyang Li: Writing – original draft. Hongpeng Zhou: Writing – original draft. Hongyu Zhang: Writing – original draft. Jixing Li: Writing – original draft. Quanying Liu: Funding acquisition, Writing – original draft. Wei Huang: Funding acquisition, Writing – original draft. Yizhuo Lu: Writing – original draft. Zijiao Chen: Writing – original draft. Jingyuan Sun: Conceptualization, Data curation, Funding acquisition, Investigation, Methodology, Project administration, Resources, Supervision, Visualization, Writing – original draft, Writing – review and editing.

## Notes

### Competing Interest Statement

The authors have declared no competing interest.

### Summary of Updates

1. Revised Figures 2, 3, and 4 and Tables 4 and 5; added Tables 6 and 7. 2. Added systematic quantitative benchmarks for different methods, and analyzed the limitations and contradictions of the studies. 3. Demonstrated the paradigm shift and the defining boundaries of the foundation model, and explored computational neuroscience theory in depth. 4. Critically discussed the merits of different metrics for evaluating brain decoding, and introduced the issue of the dual use of adversarial brain data attacks and brain decoding techniques. 5. Added new authors.

